# Automatic and fast encoding of representational uncertainty underlies probability distortion

**DOI:** 10.1101/2019.12.17.879684

**Authors:** Xiangjuan Ren, Huan Luo, Hang Zhang

## Abstract

Humans do not have an accurate representation of probability information in the environment but distort it in a surprisingly stereotyped way (“probability distortion”), as shown in a wide range of judgment and decision-making tasks. Many theories hypothesize that humans automatically compensate for the uncertainty inherent in probability information (“representational uncertainty”) and probability distortion is a consequence of uncertainty compensation. Here we examined whether and how the representational uncertainty of probability is quantified in the human brain and its relevance to probability distortion behavior. Human subjects kept tracking the relative frequency of one color of dot in a sequence of dot arrays while their brain activity was recorded by magnetoencephalography (MEG). We found converging evidence from both neural entrainment and time-resolved decoding analysis that a mathematically- derived measure of representational uncertainty is automatically computed in the brain, despite it is not explicitly required by the task. In particular, the encodings of relative frequency and its representational uncertainty respectively occur at latencies of approximately 300 ms and 400 ms. The relative strength of the brain responses to these two quantities correlates with the probability distortion behavior. The automatic and fast encoding of the representational uncertainty provides neural basis for the uncertainty compensation hypothesis of probability distortion. More generally, since representational uncertainty is closely related to confidence estimation, our findings exemplify how confidence might emerge prior to perceptual judgment.

## Introduction

Humans do not have an accurate representation of probability or relative-frequency information in the environment but distort it in a surprisingly stereotyped way. Typically, small probability is overestimated and large probability underestimated, which can be well fit by a linear-in-log-odds (LLO) model with two parameters (see [1] for a review). Such “probability distortion” phenomena occur in a variety of judgment and decisionmaking tasks such as relative-frequency estimation [2–4], confidence rating [5–7], decision under risk [8–11], and are also widely reported in animal behaviors [12–14]. However, the neural computations that accompany such probability distortions remain largely unknown.

One computation that is assumed to be central to probability distortion [9, 15–17] is compensation for the uncertainty inherent in the representation of noisy probability information (“representational uncertainty”). In particular, recent comparison among an extensive set of computational models of probability distortion [9] suggests that in their judgment and decision-making people take into account representational uncertainty that is proportional to *p*(1–*p*), where *p* denotes probability or relative frequency. That is, representational uncertainty is zero at *p* = 0 or 1 and maximal at *p* = 0.5. The *p*(1– *p*) form had also been proved by Lebreton et al. [18] as a measure of uncertainty that correlates with the explicitly reported confidence for a specific judged value. Specifically, their fMRI results showed that during the valuation process, this uncertainty measure is automatically encoded in the ventromedial prefrontal cortex (vmPFC) — the brain region known for encoding value—even in the absence of explicit confidence rating. Motivated by previous neuroimaging studies and computational modelling, we ask whether representational uncertainty of probability information can be automatically encoded in human brain and if yes, whether this encoding can precede the explicit judgment of probability, as theories of probability distortion would expect [9, 15–17].

In the present study, we designed a new experimental paradigm where a major component of the representational uncertainty of probability information (i.e. *p*(1–*p*), see Results for details) was varied continuously over time. By using time-resolved neural measurements and assessing the temporal coupling between stimulus variables and ongoing brain activities, we examined whether and how the encoding of representational uncertainty proceeds in time and relates to probability distortion.

Twenty-two human subjects participated in the study and were instructed to continuously track the relative-frequency (*p*) of one color of dots in a sequence of dot arrays (Fig 1A), during which their brain activities were recorded by magnetoencephalography (MEG). First, we found that, though *p* was the only variable that subjects needed to track while *p*(1–*p*) was task-irrelevant, the periodic changes of *p*(1–*p*) as well as *p* entrained neural rhythms, supporting an automatic tracking of representational uncertainty in the brain even when it is task-relevant. Next, by using a time-resolved decoding analysis to delineate the temporal course, we further found that the encoding of *p* and *p*(1–*p*) peaked around 300 and 400 ms, respectively. Finally, the relative strength of the neural responses to the two variables (*p* and *p*(1–*p*)) in the parietal region correlated with the variation of probability distortion behavior across individuals. Taken together, our results provide neural evidence for an automatic, fast encoding of representational uncertainty in the human brain that might underlie probability distortion observed in a wide range of human behaviors.

**Fig 1.**
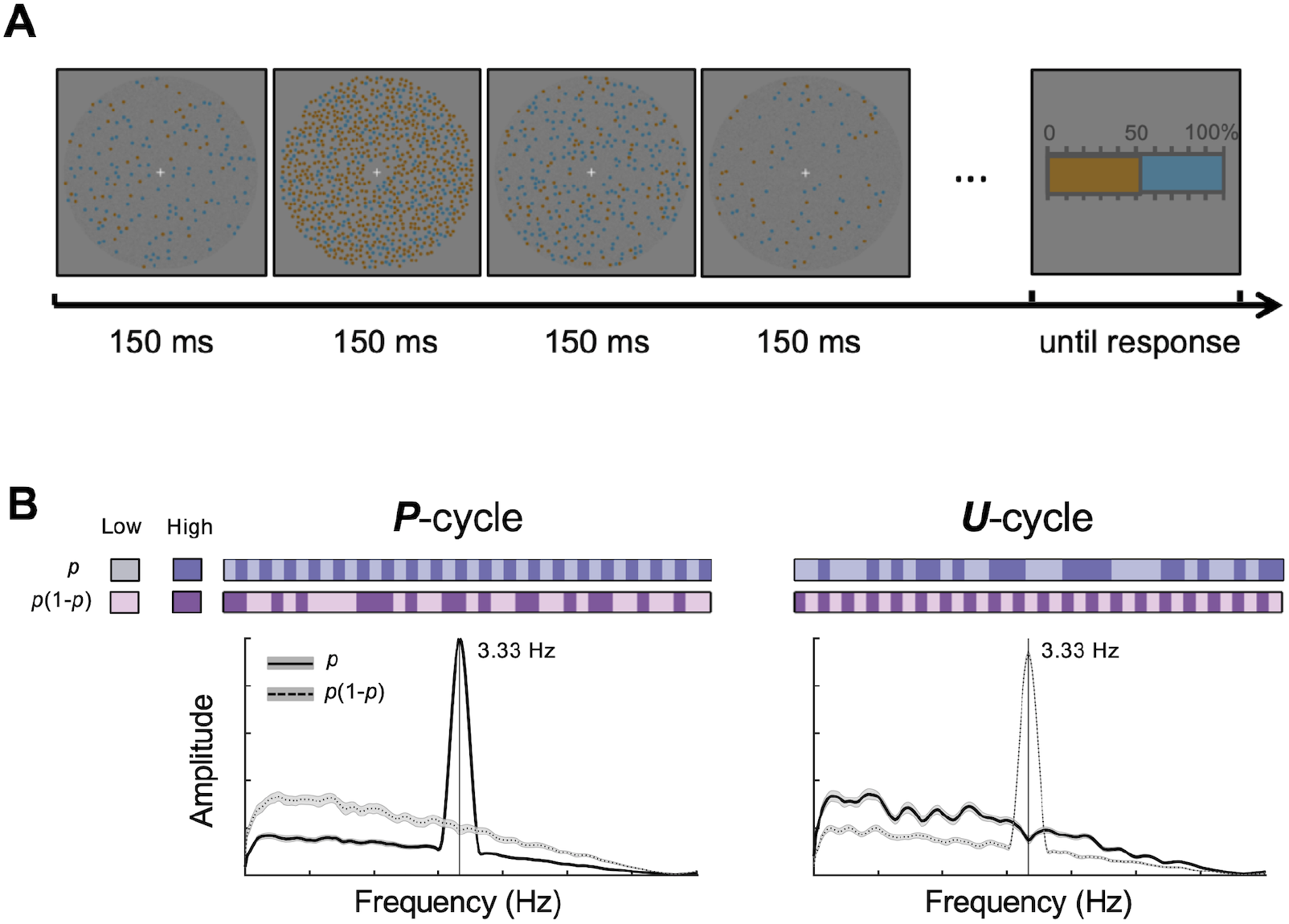
Experimental design and behavioral results. **(A)** Relative-frequency tracking and judgment task. On each trial, a sequence of orange and blue dot arrays was presented at a rate of 150 ms per display. Subjects were asked to fixate on the central fixation cross and track the relative frequency of orange (or blue) dots in each display until a 0–100% response scale appeared. Subjects then clicked on the scale to report the relative frequency in the last display. **(B)** The *p* and *p*(1–*p*) sequences in an example *P*-cycle or *U*-cycle trial. The value of *p* on each display was randomly chosen from a uniform distribution ranging from 0.1 to 0.9. In *P*-cycles (left panels), the *p* sequence was generated in such a way that the *p* value in different displays alternated between lower (<0.5) and higher (>0.5) values, resulting in a frequency spectrum peaking at 3.33 Hz (solid line), while the corresponding *p*(1–*p*) sequence was aperiodic (dashed line). In *U*-cycles (right panels), it was the reverse: the *p*(1–*p*) sequence alternated between lower (<0.21) and higher (>0.21) values in cycles of 3.33 Hz (dashed line), while the *p* sequence was aperiodic (solid line). To illustrate the periodicity or aperiodicity in the *p* and *p*(1–*p*) sequences, the lower and higher *p* values, and lower and higher *p*(1–*p*) values are coded in different discrete colors in the bars above the frequency spectrum plots. Note that this discretization is only for illustration purposes and all data analyses were still based on continuous values of *p* or *p*(1–*p*).

## Results

As shown in Fig 1A, on each trial, subjects saw a sequence of displays consisting of isoluminant orange and blue dots that was refreshed every 150 ms, and were instructed to track the relative-frequency of dots in one color (i.e. orange or blue, counterbalanced across subjects). To ensure subjects’ persistent tracking throughout the trial, the sequence of displays ended after random duration and subjects needed to report the relative-frequency of the last display by clicking on a percentage scale afterwards.

A 2 (*P*-cycle or *U*-cycle) by 2 (*N*-small or *N*-large) experimental design was used, as we describe below. Specifically, for *P*-cycle trials, the sequence of displays was generated according to the *p* value and the displays alternated between lower values of *p* (<0.5) and higher values of *p* (>0.5), with every two displays forming a cycle. As the consequence, the values of *p* in *P*-cycle trials varied at a rhythm of 3.33 Hz, while the values of *p*(1–*p*) would be aperiodic. In contrast, for *U*-cycle trials (*U* for uncertainty), the sequence of displays was generated according to the value of *p*(1–*p*) in a similar way so that the displays alternated between lower values of *p*(1–*p*) (<0.21) and higher values of *p*(1–*p*) (>0.21) in cycles of 3.33 Hz, while the values of *p* were aperiodic. In other words, in *P-cycle* trials, the *p* value underwent a rhythmic fluctuation and the *p*(1– *p*) value followed an aperiodic random course, whereas the opposite pattern occurred for *U*-cycle trials. Fig 1B illustrates the *P*-cycle (left) and *U*-cycle (right) conditions. Note that the discrete colors used in Fig 1B to code the values of *p* and *p*(1–*p*) are only for visualization purposes. The actual values of *p* and *p*(1–*p*) were randomly chosen from continuous uniform distributions (see Methods).

Our experimental design had two important features. First, the values of *p* and *p*(1–*p*) were statistically independent of each other. Second, the values of *p* and *p*(1– *p*) were dissociated in periodicity. Note that subjects’ task was always to attend to *p* (relative frequency), and *p*(1–*p*) was completely task-irrelevant in both *P*-cycle and *U*- cycle trials.

Besides, the total number of dots in a display (numerosity, denoted *N*) was not constant but varied from display to display, independently of *p* or *p*(1–*p*), ranging from 10 to 90 (*N*-small trials) or from 100 to 900 (*N*-large trials). The introduction of *N*-small versus *N*-large trials served as a means to manipulate the slope of probability distortion [1] so as to reveal the possible links between behavior and brain activity.

### Behavioral probability distortions quantified by LLO

All the 22 subjects performed well on reporting the relative-frequency of the last display, whose subjective estimate *π*(*p*) was highly correlated with the objective relative-frequency *p* (all Pearson’s *r* > 0.71, *P* < 0.001). Given that the stimulus sequence might end unexpectedly, if subjects had failed to track the sequence, they might have missed the last stimulus. Subjects’ sensible reports thus provide evidence that they had tracked the change of *p* throughout the trial as instructed.

Fig 2A shows the reported *π*(*p*) of one representative subject, who showed a typical inverted-*S*-shaped probability distortion—overestimating small *p* and underestimating large *p*. To better visualize the deviation of *π*(*p*) from *p*, we plot *π(p*)- *p* as a function of *p* for each individual subject (S1 Fig). As expected, most subjects exhibited the typical inverted-*S*-shaped probability distortion and a few subjects exhibited the opposite *S*-shaped distortion, a profile also consistent with previous findings [1, 4].

**Fig 2.**
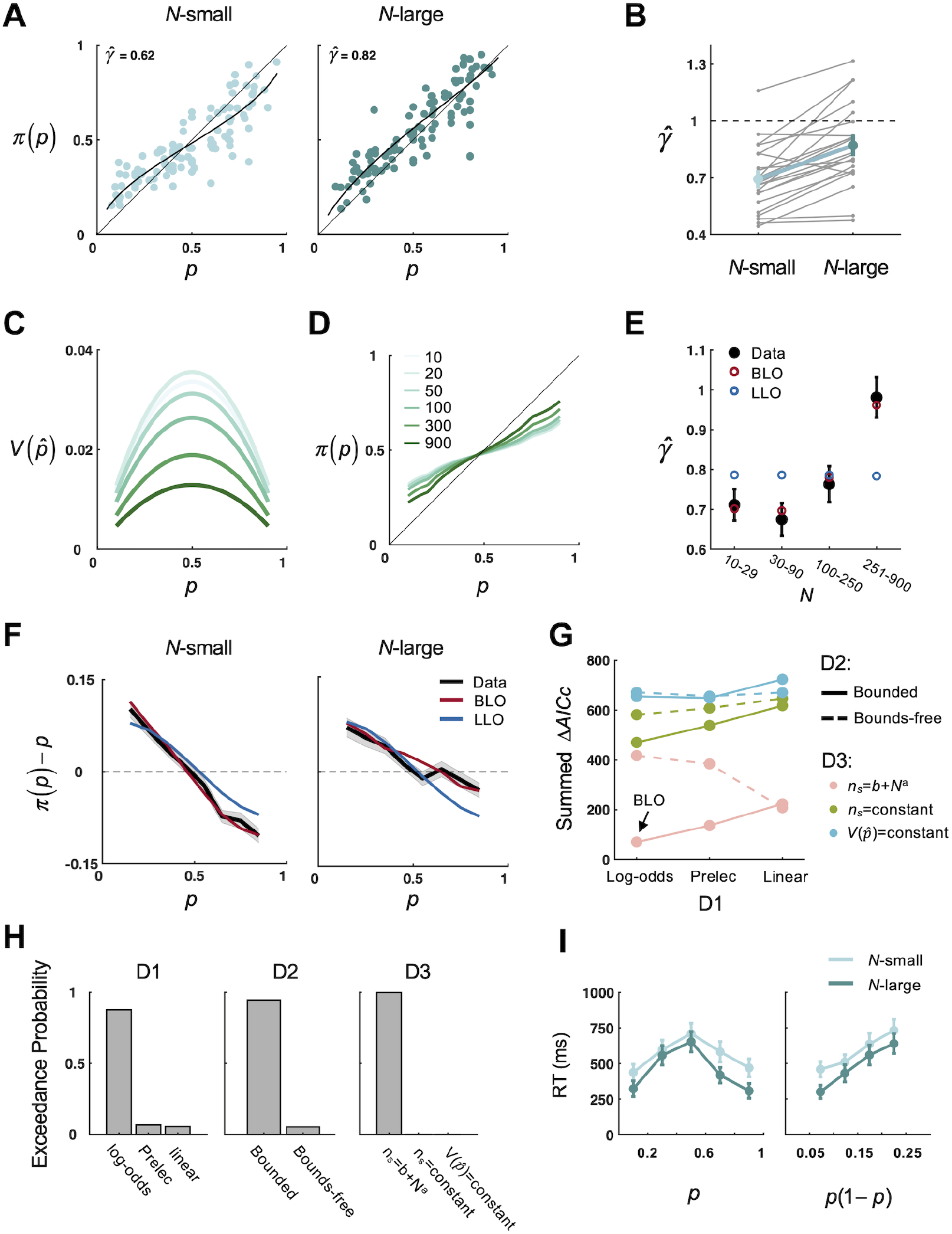
Behavioral and modeling results. **(A) Distortion of relative-frequency of one representative subject.** The subject’s reported relative-frequency *π*(*p*) is plotted against objective relative frequency *p*, separately for the *N*-small and *N*-large conditions. Each dot is for one trial. Black curves denote LLO model fits. **(B) Slope of distortion** 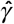 **for the *N*-small and *N*-large conditions.** Each gray line is for one subject. The thick green line denotes the group mean. Error bars denote SEM. **(C) Illustration of representational uncertainty** 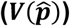) **as a function of *p* and *N*.** Larger values of *N* are coded in darker colors (same as **D**). The maximal sample size is assumed to follow the form *n_s_* = *b* + *N^α^*, with *a*=0.42 and *b*=1.91 (median estimates across subjects). **(D) Illustration of probability distortion predicted by BLO: *π(p)* as a function of *p* and *N***. Larger values of *N* are coded in darker colors. **(E) Slopes of distortion: data versus model fits.** According to the value of *N* in the last display, all trials were divided into four bins and the slope of distortion, 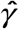, was estimated separately for each bin. Gray filled circles denote data. Error bars denote SEM. Red and blue circles respectively denote BLO and LLO model fits. **(F) π(*p*) - *p* as a function of *p* and *N***. Black curves denote smoothed data on the group level, separately for the *N*- small and *N*-large conditions. Shadings denote SEM. Red and blue curves respectively denote BLO and LLO model fits. **(G) Results of factorial model comparison: summed** Δ**AICc**. Lower values of ΔAICc indicate better fits. The BLO model outperformed all the alternative models. **(H) Protected exceedance probability on each model dimension.** Each panel is for model comparison on one dimension. Each assumption of the BLO model outperformed the alternative assumptions on its dimension. **(I) Response time as an additional index for representational uncertainty.** The mean response time (RT) for subjects to initiate their relative-frequency report is plotted against binned *p* or *p*(1–*p*), separately for the *N*-small and *N*-large conditions. Error bars denote SEM.

We next used the LLO model to quantify such inverted-*S*- or *S*-shaped distortions, summarizing each subject’s probability distortion behavior for each trial type (*P*-cycle or *U*-cycle) and dot numerosity (*N*-small or *N*-large) conditions by two parameters fitted from *π*(*p*) (see Methods): slope 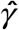 and crossover point 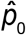. According to 2 (*P*-cycle or *U*-cycle) by 2 (*N*-small or *N*-large) repeated-measures ANOVAs separately performed for the estimated 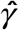 and 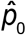, numerosity showed a significant influence on 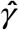 (*F*(1, 63) = 57.82, *P* < 0.001, 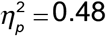) and a statistically significant but small influence on 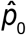 (*F*(1, 63)=4.24, *P* = 0.044, 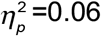). No other main effects or interactions reached significance (all *P* > 0.26). In particular, the 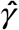 estimated from *N*-large trials was greater than that of *N*-small trials by 25.75% (Fig 2B). Thus our subsequent analysis would mainly focus on 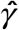 and collapse the two cycle conditions to estimate 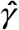. As illustrated in Fig 2A, a greater 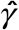 implies a less curved inverted-*S*- shaped distortion (for 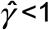) or a more curved *S*-shaped distortion (for 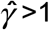).

It is noteworthy that LLO was only used as a way to quantify probability distortions and their differences between conditions. The LLO model by itself could not explain why the slope of distortion would differ between numerosity conditions, as observed here (Fig 2AB). In contrast, the observed different 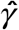 between the *N*-small and *N*- large conditions could be well captured by the bounded log-odds (BLO) model [9], which compensates for the *p*(1 –*p*) form of representational uncertainty, with greater 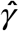 implying lower representational uncertainty. Next we will present the BLO model and how it may account for our behavioral results.

### Bounded Log-Odds model and its behavioral evidence

Zhang, Ren, and Maloney [9] proposed the BLO model as a rational account of probability distortion, which can explain why probability distortion may vary with task or individual. BLO has three main assumptions: (1) probability is internally represented as log-odds, (2) representation is truncated to a bounded scale, and (3) representational uncertainty is compensated in the final estimate of probability, so that estimates associated with higher uncertainty will be discounted to a greater extent. See S1 Text for details. A full description of BLO would be out of the scope of the present article and below we only focus on the uncertainty compensation assumption.

According to BLO, when representational uncertainty (denoted 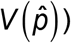 is higher, the subjective estimate of probability is less contributed by the percept of the objective probability and more by a prior estimate, which implies that the higher the representational uncertainty, the shallower the slope of probability distortion. As illustrated in Fig 2C, the representational uncertainty modeled in our experiment (see S1 Text) is proportional to *p*(1–*p*), that is, an inverted-*U*-shaped function of *p*. Meanwhile, for a median subject, 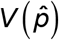 first slightly increases with *N* (Fig 2C, from *N*=10 to *N*=20) and then dramatically decreases with *N* (Fig 2C, from *N*=20 to *N*=900). Consequently, BLO predicts that the slope of distortion 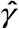 is not necessarily constant, but may slightly decrease with *N* for very small *N* and mostly increase with *N* (Fig 2D), which is indeed observed in our experiment (Fig 2B & 2E).

We are aware that representational uncertainty 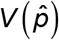 here is not equivalent but proportional to *p*(1–*p*). In the neural analysis we will present, we chose to focus on the encoding of *p*(1–*p*) (as a proxy for representational uncertainty) instead of 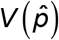, because 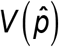 was highly correlated with numerosity *N* and thus would be difficult to be separated from the latter in brain activities. In contrast, *p*(1–*p*) and *N* were independent of each other by design. Besides, *p*(1–*p*) has a nice connection with the “second-order valuation” proposed by Lebreton et al. [18].

#### Model versus data

Zhang et al. [9] has provided behavioral evidence for the BLO model in two different tasks including relative-frequency judgment. Here we performed similar tests on our behavioral data and fit the BLO model to the reported *π*(*p*) for each subject using maximum likelihood estimates. The performance of the LLO model served as a baseline. When evaluating LLO’s performance in fitting the data, we estimated one set of parameters for all conditions instead of estimating different parameters for different conditions as when we applied LLO as a measuring tool for probability distortions. In Fig 2F, we plot the observed *π*(*p*)- *p* (smoothed and averaged across subjects) as a function of *p* separately for different numerosity conditions and contrasted it with the BLO and LLO model predictions. The BLO prediction agreed well with the observed probability distortion function, while LLO largely failed. The advantage of BLO over LLO was even more pronounced in plots for individual subjects (S1 Fig).

We further examined how well BLO could predict the observed numerosity effect in the slope of distortion. For each subject, we divided the trials of each numerosity condition evenly into two bins according to the value of *N* in the last display (i.e., the display that was reported) and then estimated 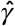 for each of the four bins. The pattern of 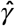 in real data—initial slight decrease and subsequent dramatic increase with *N*— was quantitatively predicted by the fitted BLO model (Fig 2E).

#### Factorial model comparison

Similar to Zhang et al. [9], we performed a factorial model comparison [19] to test whether each of the three assumptions in the BLO model outperformed plausible alternative assumptions in fitting behavioral data. Accordingly we constructed 3 *×* 3 *×* 2 = 18 models and fit each model for each subject using maximum likelihood estimates (see S2 Text for details). The Akaike information criterion with a correction for sample sizes, AICc [20, 21], was used as the metric of goodness-of-fit. Lower AICc indicates better fit. For each subject, the model with lowest AICc was used as a reference to compute ΔAICc for each model. According to the summed ΔAICc across subjects, BLO was the best model among the 18 models (Fig 2G).

We also evaluated whether each of BLO’s three assumptions was the best among its alternatives on the same dimension. In particular, we used the group-level Bayesian model selection [22–24] to compute the probability that each specific model outperforms the other models in the model set (“protected exceedance probability”) and marginalized the protected exceedance probabilities for each dimension (i.e., adding up the protected exceedance probabilities across the other two dimensions). Indeed, all three assumptions of the BLO model in the present study—log-odd, bounded, and compensation for 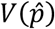 (with *n_s_* = *b* + *N^a^*)—outperformed their alternatives with probabilities of 87.7%, 94.6% and 99.8%, respectively (Fig 2H).

Furthermore, we performed a model recovery analysis (similar to [25]) to confirm that the advantage of BLO in factorial model comparison is real and does not result from model mis-identification. In particular, we generated 50 synthetic datasets for each of the 18 models. All the datasets generated from BLO were best fit by BLO. Out of the 850 datasets generated from the other models, only 0.24% were mis-identified to BLO (see S2A Fig for details).

We also verified that the parameters of BLO could be reasonably well identified (S2B Fig). Even if some of the BLO parameters were not perfectly identified, it would not influence the results of the neural analysis we report below, most of which did not rely on the estimated BLO parameters.

### Evidence for representational uncertainty in response time

The above modeling analysis on subjects’ reported relative frequency (i.e. model fits as well as the factorial model comparison) suggests that subjects might compensate for representational uncertainty that is proportional to *p*(1–*p*) and the *N*-large condition was associated with lower uncertainty than the *N*-small condition.

The response time (RT) of reporting relative frequency (defined as the interval between the response screen onset and the first mouse move) provides an additional index for representational uncertainty, given that lower uncertainty would lead to shorter RTs. We divided the values of *p* into five bins and computed RT for each bin and separately for the *N*-small and *N*-large trials. According to a 5 (*p* bins) by 2 (*N*- small vs. *N*-large) repeated-measures ANOVA on RTs, both the main effects of the *p* value (*F*(4, 189) = 29.02, *P* < 0.001, 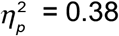) and numerosity (*F*(1, 189) = 25.61, *P* < 0.001, 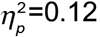) were significant. Consistent with Lebreton et al. [18], RTs were longest at *p*=0.5 and shortest when *p* was close to 0 or 1 (Fig 2I, left). More precisely, RTs increased almost linearly with *p*(1–*p*) (Fig 2I, right), in agreement with what one would expect if representational uncertainty is proportional to *p*(1–*p*). Besides, RTs were shorter in *N*-large trials than in *N*-small trials, which echoes the lower 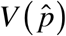 for larger *N* (Fig 2C) and provides further evidence that the *N*-large condition was accompanied by lower representational uncertainty.

In sum, we found that a considerable portion of variability in subjects’ probability distortion functions and RT patterns can be accounted by differences in representational uncertainty. One may wonder whether the effects of representational uncertainty can be explained away by task difficulty. We doubt not, because higher representational uncertainty does not necessarily correspond to higher task difficulty. For example, representational uncertainty for relative-frequency estimation is proportional to *p*(1–*p*), which is maximal at *p*=0.5 and minimal when *p* is close to 0 or 1. In contrast, regarding difficulty, there seems to be little reason to expect that relative-frequency estimation should be more difficult for *p*=1/2 than for *p*=1/3, or be more difficult for *p*=1/3 than for *p*=1/4. As another counter-example, representational uncertainty modeled in the present study is higher in the *N*-small condition than in the *N*-large condition (Fig 2C), but there seems to be little reason to expect relative-frequency estimation to be more difficult for displays with fewer dots.

### Neural entrainment to periodic changes of *p* or *p*(1–*p*)

After showing that the BLO models can well capture the behavioral results, we next examined whether the brain response could track the periodically changing *p* values (*P*-cycle) or *p*(1–*p*) values (*U*-cycle) in the stimulus sequence. Recall that the *P*-cycle and *U*-cycle trials had identical individual displays and differed only in the ordering of the displays: In *P*-cycle trials, *p* alternated between small and large values in cycles of 3.33 Hz while *p*(1–*p*) was aperiodic; in *U*-cycle trials, *p*(1–*p*) alternated in cycles of 3.33 Hz while *p* was aperiodic.

As shown in Fig 3, the brain response indeed tracked the periodic changes of *p* and *p*(1–*p*). In particular, in *P*-cycle trials (Fig 3, left), the phase coherence between the periodic *p* values and the MEG time series reached significance at 3.33 Hz (permutation test, *FDR* corrected *P_FDR_* < 0.05), mainly in the occipital and parietal sensors. Importantly, significant phase coherence was also found at 3.33 Hz in *U*-cycle trials between the periodic *p*(1–*p*) values and the MEG time series (Fig 3, right). That is, the brain activity was also entrained to the periodic changes of the representational uncertainty of *p* (i.e. *p*(1–*p*)). Given that subjects were only asked to track the value of *p* but not *p*(1–*p*), the observed neural entrainment to the task-irrelevant *p*(1–*p*) suggests an automatic encoding of representational uncertainty in the brain.

**Fig 3.**
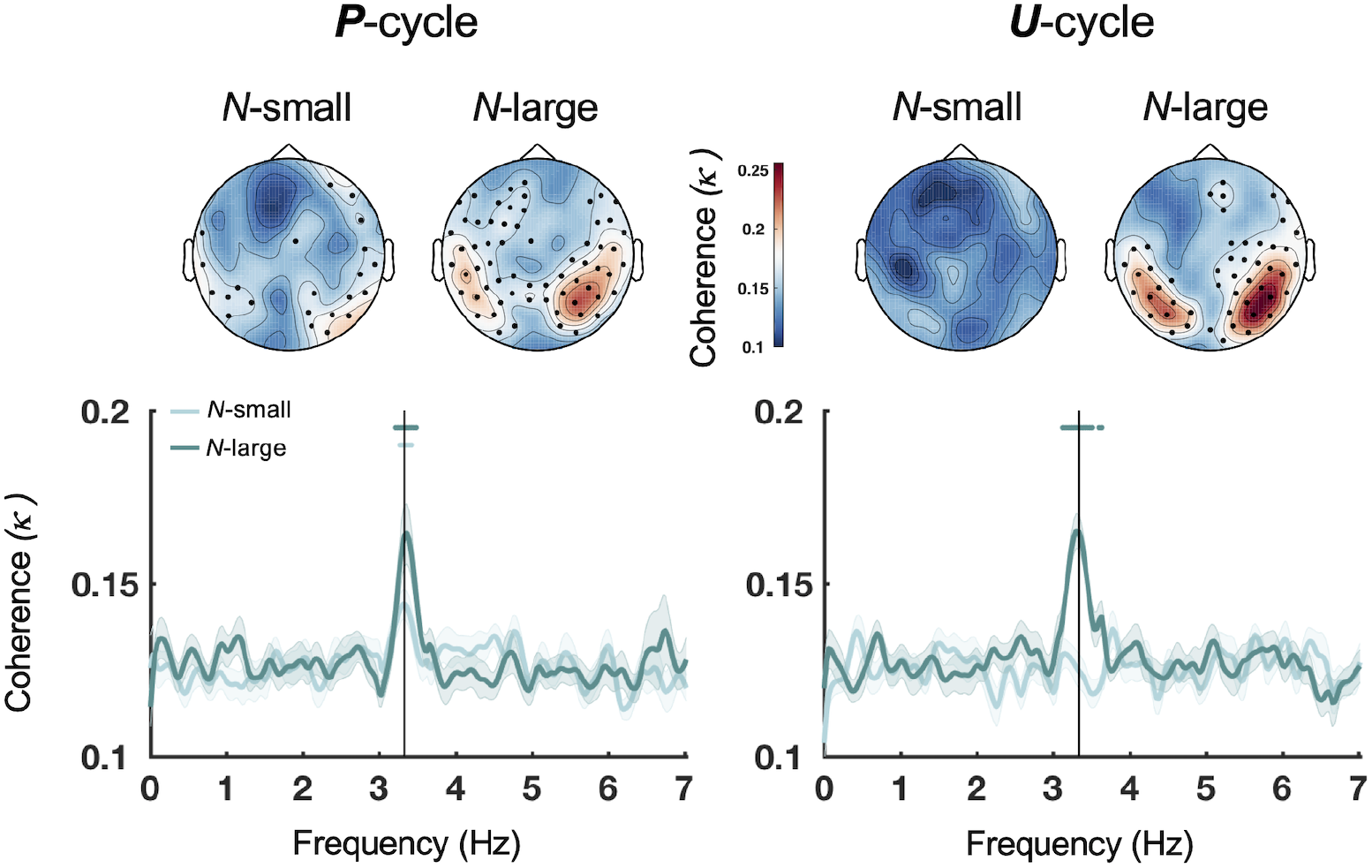
Results of phase coherence analysis. Grand-averaged phase coherence spectrum for magnetometers. **Left**: The phase coherence (*κ*) between the periodic *p* values and MEG time series in *P*-cycle trials. **Right**: The phase coherence between the periodic *p*(1–*p*) values and MEG time series in *U*-cycle trials. Light and dark green curves respectively denote the *N*-small and *N*-large conditions. Shadings denote SEM across subjects. The vertical line marks 3.33 Hz. Dots above the spectrum mark the frequency bins whose phase coherence was significantly above chance level (permutation tests, *P_FDR_* <0.05 with *FDR* correction across frequency bins). **Insets:** Grand-averaged phase coherence topography at 3.33 Hz for each cycle and numerosity condition. Solid black dots denote sensors whose phase coherence at 3.33 Hz was significantly above chance level (permutation tests, *P_FDR_*<0.05 with *FDR* corrected across magnetometers).

Notably, the significant phase coherences between the periodic *p* or *p*(1–*p*) values and MEG time series were not simply because the MEG time series consisted of 3.33 Hz frequency components. In a control analysis, we calculated the phase coherence between the aperiodic variable (i.e. *p* in *U*-cycle trials or *p*(1–*p*) in *P*-cycle trials) and the same MEG time series, and found that there were no significant peaks at 3.33 Hz (S5B Fig). Moreover, we had carefully controlled potential confounding variables in our experimental design. For example, *p* and *p*(1–*p*) were linearly independent of each other, both of which had negligible correlations with numerosity or low-level visual features such as luminance, contrast, and color variance (S5A Fig). Furthermore, different from *p* and *p*(1–*p*), these potential confounding variables were associated with similar entrainment responses in *P*-cycles and *U*-cycles (S5B Fig).

### Behavioral probability distortion and the neural entrainment to *p* and *p*(1– *p*)

We next examined the relationship between behavioral probability distortion and the neural entrainment to *p* and *p*(1–*p*) on a subject-by-subject basis. Specifically, the phase coherence between a specific variable (*p* or *p*(1–*p*)) and the MEG time series can be considered as a measure of the strength of neural responses to the variable [26]. We defined

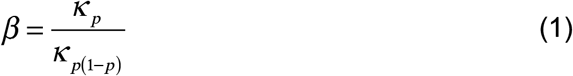

to quantify the relative strength of *p* to *p*(1–*p*) in neural responses, where *κ_p_* denotes the phase coherence for *p* in *P*-cycle trials, and *κ*_*p*(1-*p*)_ denotes the phase coherence for *p*(1–*p*) in *U*-cycle trials. A higher value of *β* would imply a stronger neural encoding of relative-frequency or a weaker encoding of representational uncertainty and is thus supposed to yield probability distortions of a greater slope [9]. Note that both the behavioral and neural measures we defined below were intra-subject ratios that were unitless and scale free, thus not subject to the potential scaling issues in inter-individual correlation [27].

Given that the estimated slope of distortion 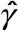 was greater in *N*-large trials than in *N*-small trials, we would expect *β* to change in the same direction across the numerosity conditions. That is, suppose we define 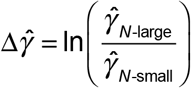 and 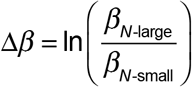, there should be a positive correlation between 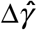 and Δ*β*.

As shown in Fig 4, there was indeed a significant correlation between behavioral and neural measures across subjects in the parietal region (Pearson’s *r* = 0.67, *FDR*- corrected one-tailed *P_FDR_* = 0.033), associating the neural entrainment to *p* and *p*(1–*p*) with probability distortion behaviors.

**Fig 4.**
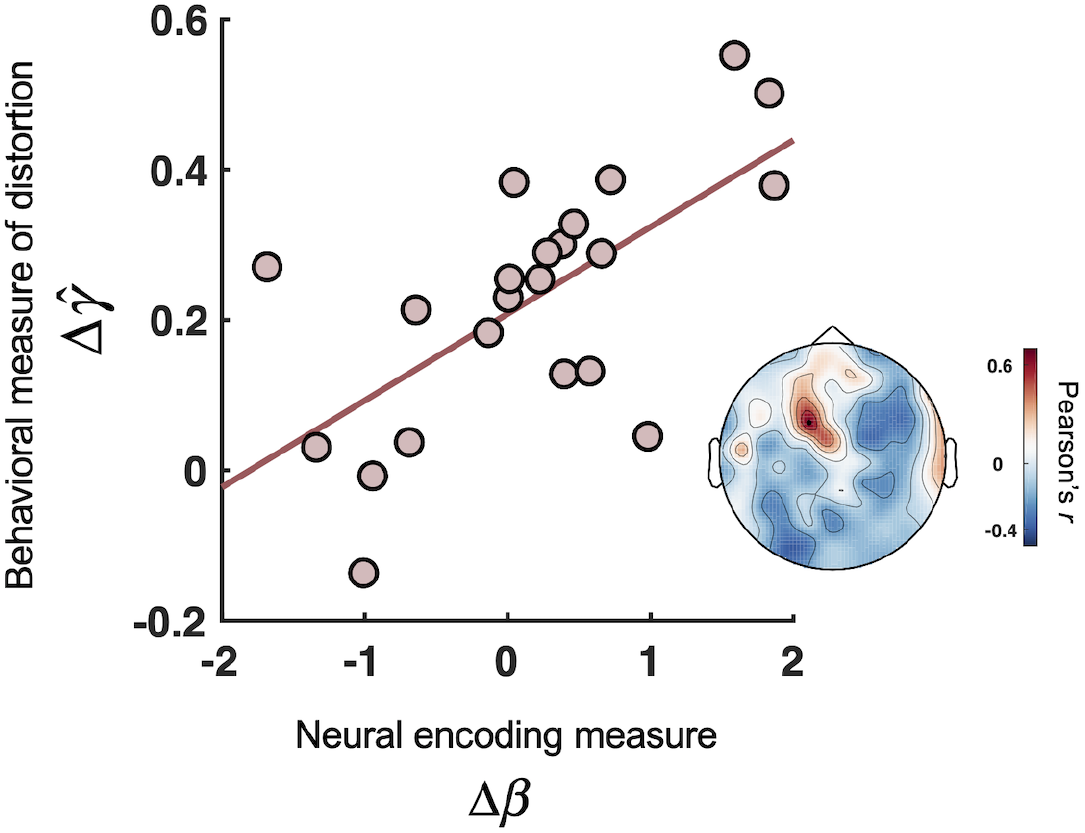
Neural responses to *p* and *p*(1–*p*) predict the slope of probability distortion. We defined 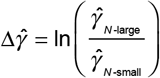 and 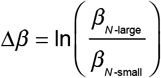 to respectively quantify how much the behavioral measure of the slope of distortion, 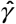, and the relative strength of *p* to *p*(1–*p*) in neural responses, *β*, changed across the two numerosity conditions. **Main plot**: Correlation between 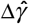 and the Δ*β* at the parietal magnetometer channel MEG0421. The logarithm transformation was used only for visualization. Each dot denotes one subject. **Inset**: Correlation coefficient topography between 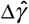 and Δ*β*, on which MEG0421 is marked by a solid black dot.

### Neural encoding of *p* and *p*(1–*p*) over time and the associated brain regions

After establishing the automatic tracking of cyclic changes in *p*(1–*p*) and its behavioral relevance, we next aimed to delineate the temporal dynamics of *p* and *p*(1–*p*) in the brain signals. Recall that on each trial one of the two variables of our interest (*p* or *p*(1– *p*)) changed periodically and the other was aperiodic. Therefore, we could decode the temporal course of aperiodic *p*(1–*p*) in *P*-cycles trials and aperiodic *p* in *U*-cycle trials.

In particular, we performed a time-resolved decoding analysis based on all 306 sensors (see Methods) using a regression approach that has been employed in previous EEG and MEG studies [28, 29] including ours [30–32]. The intuition of the time-resolved decoding analysis is as follows. Suppose the onset of each new *p* or *p*(1–*p*) value in the stimulus sequence evokes phase-locked brain responses that extend across time and whose magnitudes are proportional to the encoded value. The resulting MEG time series would then be a superposition of the responses to all the preceding stimuli in the sequence. But as soon as the values of an encoded variable are not correlated over time (i.e. free of autocorrelation), their brain response profiles are separable. In particular, for any stimulus in the sequence, we can use the MEG signals at a specific delay after the stimulus onset to predict the value of the stimulus. Importantly, different from the phase coherence analysis (Fig 3) that only reflects the overall strength of the brain responses, this time-resolved decoding analysis allows us to assess how *p* and *p*(1–*p*) are encoded over time.

Fig 5 plots the temporal course of the decoding performance for *p* (left) and *p*(1– *p*) (right). We found that both *p* and *p*(1–*p*) were successfully decoded from the MEG signals in *N*-large trials (cluster-based permutation test, *P_cluster_* < 0.05). In particular, the decoding performance for *p* peaked around 327 ms after stimulus onset (Fig 5, left), while that for *p*(1–*p*) peaked around 419 ms (Fig 5, right). In other words, the neural encodings of relative frequency and its representational uncertainty had distinct time courses, with the latter occurring approximately 100 ms later than the former (also see S8 Fig for the decoding performance of individual subjects).

**Fig 5.**
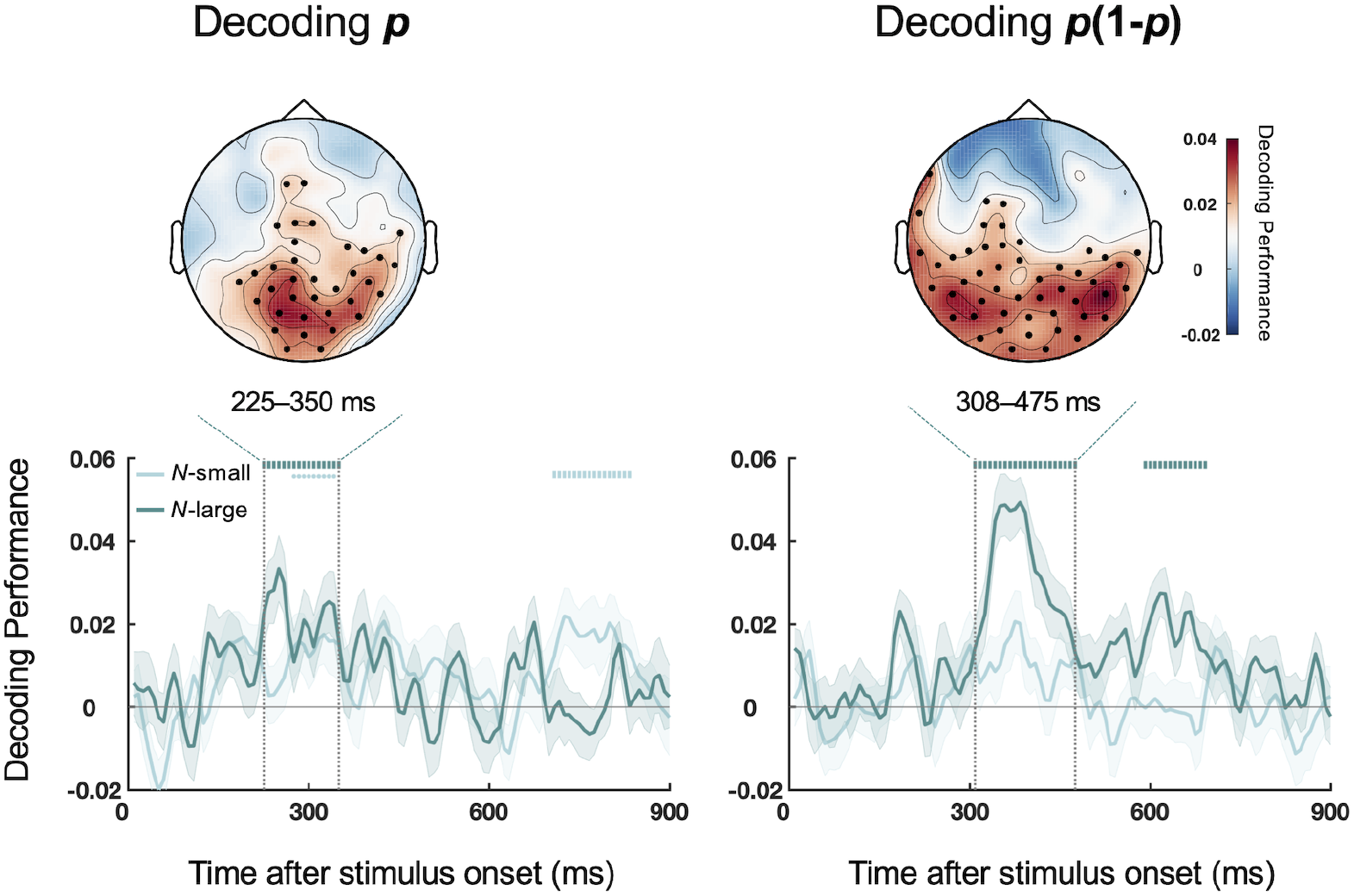
Results of decoding analyses. **Main plots**: Time-resolved decoding performance over different time lags for *p* (left panel) and *p*(1–*p*) (right panel), separately for the *N*-small (light green) and *N*-large (dark green) conditions. Shadings denote SEM across subjects. Symbols above the curves indicate time lags that had above-chance decoding performance (cluster-based permutation tests), with vertical bars representing *P_cluster_*<0.01 and dots representing 0.01≤*P_cluster_*<0.05. The decoding performance for *p* peaked at around 327 ms after stimulus onset, while that for *p*(1–*p*) peaked at around 419 ms. **Insets**: Topography of spatial decoding performance for the time window (highlighted by the funnel-shaped dashed lines) that contained the peak decoding performance in the time-resolved decoding analysis. Solid black dots indicate the sensor locations with above-chance decoding performance (*P_FDR_* < 0.05) for the *N*- large condition. (Spatial decoding in the same time windows for the *N*-small condition resulted in no statistically significant sensors.) The encodings of *p* and *p*(1–*p*) involved overlapping parietal-occipital regions, including the parietal region where Δ*β* and 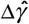 were positively correlated (Fig 4).

In contrast, none of the potential confounding variables (numerosity, luminance, etc.) showed temporal courses similar to *p* or *p*(1–*p*) when the same time-resolved decoding procedure was applied (S5C Fig). The automatic encoding of the taskrelevant *p*(1–*p*) might seem surprising. To further verify that this was not an artifact of potential low-level confounds, we performed an additional decoding analysis for *p*(1– *p*) based on the cross-validated version of confound regression (CVCR) [33], where the confounding variables were regressed out from MEG time series before time- resolved decoding analysis (see S7 Text for details). The decoded temporal course for *p*(1–*p*) was little changed (S9A Fig). A similar decoding analysis for the complete form of representational uncertainty 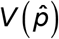 resulted in similar temporal course as that of *p*(1–*p*) (S9B Fig). In sum, we found automatic encoding of *p*(1–*p*) in brain signals even when it was task-irrelevant and when low-level confounding factors were excluded.

Further, based on the time windows that achieved the highest time-resolved decoding performance, we performed a similar decoding analysis at each sensor location (including one magnetometer and two orthogonal planar gradiometers) separately for *p* and *p*(1–*p*) to assess the associated brain regions (see Methods). We found that the encoding of the two variables involved overlapping parietal-occipital regions (Fig 5, see S7 Fig for the results of the other above-chance time windows), including the parietal region where Δ*β* and 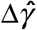 were positively correlated (Fig 4).

## Discussion

We used MEG to investigate probability distortions in a relative-frequency estimation task and found that the human brain encodes not only the task-relevant relative frequency but also its task-irrelevant representational uncertainty. The neural encoding of the representational uncertainty occurs at as early as approximately 400 ms. These results suggest that the human brain automatically and quickly quantifies the uncertainty inherent in probability information. Our findings provide neural evidence for the theoretical hypothesis that probability distortion is related to representational uncertainty. More generally, these findings may connect to the functional role of confidence (estimation of uncertainty) during judgment and decision making.

### Neural computations underlying probability distortions

Humans show highly similar probability distortions in tasks involving probability or relative frequency, with the subjective probability typically an inverted-*S*-shaped function of the objective probability [1]. Why do people distort probability information in such a systematic way? Though inverted-*S*-shaped distortions of probability had been found in the activity of a few brain regions [34, 35], the neural computations involved in transforming objective probabilities to subjective probabilities were largely unknown. Several theories [9, 15–17] rationalize probability distortion as a consequence of compensating for representational uncertainty, in accordance with the framework of Bayesian Decision Theory [36]. In brief, the brain will discount an internal representation of probability according to the level of uncertainty so that it can achieve a more reliable estimate of the objective probability. The higher the uncertainty associated with a representation, to a greater extent the representation is discounted. By modeling human subjects’ probability distortion behavior in two representative tasks—relative-frequency estimation and decision under risk—Zhang et al. [9] found evidence that representational uncertainty proportional to *p*(1–*p*) is compensated in human representation of probability or relative frequency.

What we found in the present study is neural evidence for the encoding of *p*(1–*p*) during the evaluation of *p*. For the first time, we have shown that representational uncertainty *p*(1–*p*) is not just virtual quantities assumed in computational models of probability distortions but is really computed in the brain.

### Representational uncertainty and confidence

The representational uncertainty that concerns us lies in the internal representation of probability or relative-frequency. One concept that may occasionally coincide with such *epistemological uncertainty* but is entirely different is outcome uncertainty, which is widely studied in the literature of decision under risk [37, 38], prediction error [37] and surprise [39, 40]. For example, for a two-outcome gamble with probability *p* to receive *x1* and probability *1–p* to receive *x2*, the variance of the outcomes is proportional to *p(1–p*). In the present study, the relative-frequency estimation task we chose to investigate does not involve similar binary uncertain outcomes and thus avoids the possible confusion of outcome uncertainty with representational uncertainty. Therefore, the neural encoding of *p*(1–*p*) we found here cannot be attributed to the perception of outcome uncertainty.

In fact, representational uncertainty is more closely related to confidence, except that it is not necessarily bound to any specific judgment or decision [41]. Specifically, the encoding of representational uncertainty may be understood as a special case of the “second-order valuation” proposed by Lebreton et al. [18]. As described earlier, Lebreton et al. [18] found a principled relationship between an overt numerical judgment and the individual’s confidence about the judgment, with the latter being a quadratic function of the former. They showed in fMRI studies that such quadratic form—as a proxy to confidence—is automatically encoded in vmPFC, even when confidence rating is not explicitly required of. Using the same quadratic form as the proxy, here we found automatic encoding of the representational uncertainty of relative frequency during the tracking of a visual sequence. Meanwhile, our findings extend previous work and contribute to the confidence literature in the following three aspects.

First, we found automatic encoding of representational uncertainty even in the absence of overt judgment, not just in the absence of overt confidence estimation. In our experiment, subjects were asked to track the relative frequency in the stimulus sequence but made no overt judgment except for the last display. Moreover, displays were refreshed every 150 ms, apparently leaving no time for deliberation.

Second, we resolved the temporal course of the encoding of representational uncertainty, which would be inaccessible under the low temporal resolution of fMRI. In particular, representational uncertainty is encoded as early as 400 ms upon stimulus onset, approximately 100 ms after the encoding of relative frequency itself. The occipital-parietal region is involved in the processing of both relative-frequency and its representational uncertainty but during different time windows. The fast encoding of representational uncertainty we found echoes recent psychophysiological findings in humans [42–44] and non-human primates [45] that the confidence encoding for perceptual judgments can be detected in the brain far before confidence rating and even prior to the overt judgment or decision making.

Third, our findings connect to the functional importance of confidence encoding in information processing [41, 46, 47] that is still not well understood but receives growing attention. In particular, here we ask how the neural encoding of representational uncertainty may relate to probability distortion. By comparing experimental conditions under which the same individuals’ distortions of relative frequency differed, we found that a relatively stronger response to representational uncertainty in the parietal region corresponds to a shallower slope of distortion. We conjecture that the automatic, fast encoding of representational uncertainty might not just be *post hoc* evaluation but indeed regulate probability distortion. Meanwhile, we are aware that our current findings are based on correlational analyses and whether there is causality between the neural encoding of representational uncertainty and probability distortion still awaits future empirical tests.

### Methodology implications

The “steady-state responses” (SSR) technique [48]—using rapid periodic stimulus sequence to entrain brain responses—has been widely used with EEG/MEG to investigate low-level perceptual processes, which increases the signal-to-noise ratio of detecting the automatic brain responses to the stimuli by sacrificing temporal information. In our experiment, we constructed periodic *p* or *p*(1–*p*) sequences and showed that SSR can also be used to reveal the brain’s automatic responses to these more abstract variables. Moreover, we demonstrate the feasibility to perform time-resolved decoding for the other aperiodic variable embedded in the same sequence, whereby exploiting the advantages of both the SSR and time-resolved decoding techniques. Such design would be useful for a broad range of problems that need to dissociate the processing of two or more variables in brain activities.

## Materials and Methods

### Ethics Statement

The study had been approved by the Institutional Review Board of School of Psychological and Cognitive Sciences at Peking University (#2015-03-13c). Subjects provided written informed consent in accordance with the Declaration of Helsinki.

### Subjects

Twenty-two human subjects (aged 18–27, 13 female) participated. All of them had normal or corrected-to-normal vision and passed the Farnsworth-Munsell 100 Hue Color Vision Test [49].

### Experimental Procedures

#### Apparatus

Subjects were seated approximately 86 cm in front of a projection screen (Panasonic PT-DS12KE: 49.6 × 37.2 cm, 1024 × 768 pixels, 60-Hz refresh rate) inside the magnetically shielded room. Stimuli were controlled by a Dell computer using Matlab and PsychToolbox-3 [50, 51]. Subjects’ behavioral responses were recorded by a MEG-compatible mouse system (FOM-2B-10B from Nata technologies) and their brain activities by a 306-channel MEG system (see MEG Acquisition and Preprocessing for details).

#### Task

Each trial started with a white fixation cross on a blank screen for 600 ms, following which a sequence of displays of orange and blue dots was presented on a gray background at a rate of 150 ms per display (Fig 1A). Subjects were asked to fixate on the central fixation cross and track the relative-frequency of each display. After the sequence ended for 1000 ms, a horizontal scale (0 to 100%) appeared on the screen and subjects were required to click on the scale to indicate the relative-frequency of orange (or blue) dots on the last display. Half of the subjects reported relative frequency for orange dots and half for blue dots.

To encourage subjects to pay attention to each display, one out of six trials were catch trials whose duration followed a truncated exponential distribution (1–6 s, mean 3 s), such that almost each display could be the last display. The duration of formal trials was 6 or 6.15 s. Only the formal trials were submitted to behavioral and MEG analyses.

On each display, all dots were randomly scattered without overlapping within an invisible circle that subtended a visual angle of 12°. The visual angle of each dot was 0.2° and the center-to-center distance between any two dots was at least 0.12°. The two colors of the dots were isoluminant (CIE_orange_=[43, 19.06, 52.33], CIE_blue_=[43, – 15.49, −23.72]). Isoluminant dark gray pixels (CIE_gray_=[43, 0, 0]) were filled on the gray background (CIE_background_=[56.5, 0, 0]) between the dots as needed to guarantee each display had equal overall luminance, which prevented luminance from being a confounding factor for any abstract variables of interest.

#### Design

We adopted a steady-state response (SSR) design, which could achieve a higher signal-to-noise ratio than the conventional event-related design [48]. The basic idea was to vary the value of a variable periodically at a specific temporal frequency and to observe the brain activities at the same frequency as an idiosyncratic response to the variable. The variables of most interest in the present study were relative frequency *p* and its representational uncertainty quantified by *p*(1–*p*).

For half of the trials (referred as the *P*-cycle condition), the value of *p* in the sequence was chosen from uniform distributions with the ranges alternatively being (0.1, 0.5) and (0.5, 0.9) so that the sequence of *p* formed cycles of 3.33 Hz. For the other half trials (referred as the *U*-cycle condition), the value of *p* was chosen alternatively from (0.1 0.3) U (0.7, 0.9) and (0.3, 0.7) so that the sequence of *p*(1–*p*) (i.e. proxy for uncertainty) was in cycles of 3.33 Hz. That is, the two cycle conditions had matched individual displays but differed in which variable formed a periodic sequence. Accordingly, the *p*(1–*p*) sequence in the *P*-cycle condition and the *p* sequence in the *U*-cycle condition were aperiodic, which had little autocorrelations (S3 Text & S4 Fig).

The total number of dots on a display (numerosity, denoted *N*) was varied across trials as well as across individual displays of the same trial. The value of *N* for a specific display was independent of the value of *p* and generated as a linear transformation of a beta random variable following Beta(0.1, 10) so that the number of dots in each color (*pN* or (1–*p*)*N*), a possible confounding variable of *p*, would only have moderate correlations (Pearson’s |*r*| < 0.5) with *p*. The linear transformation was chosen to locate *N* to the range of [10, 90] for half of the trials (the *N*-small condition) and to [100, 900] for the other half (the *N*-large condition).

The numbers of dots in each color was *pN* or (1–*p*)*N* rounded to the nearest integer. As the result of rounding, the actual relative-frequency was slightly different from the originally chosen *p*. In data analysis, we would use *p* to denote the actual relative-frequency that was presented to subjects.

To summarize, there were 2 (*P*-cycle vs. *U*-cycle) by 2 (*N*-small vs. *N*-large) experimental conditions, with all conditions interleaved and each condition repeated for 6 trials in each block. Each subject completed 10 blocks of 24 trials, resulting in 50 formal trials and 10 catch trials for each condition. Before the formal experiment, there were 32 practice trials (the first 20 trials consisted of one display and the following 12 trials had the same settings as the formal experiment) for subjects to be familiar with the procedure. No feedback was available during the experiment.

### Behavioral Analysis

#### Measures of probability distortion

According to Zhang and Maloney [1], inverted- *S*- or *S*-shaped probability distortions can be well captured by the LLO model

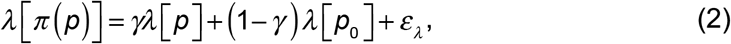

where *p* and *π*(*p*) respectively denote the objective and subjective probability or relative-frequency, 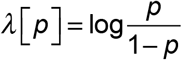 is the log-odds transformation, and *ε_λ_* is Gaussian error on the log-odds scale with mean 0 and variance 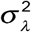. The *γ*, *p*_0_, and *σ_λ_* are free parameters. The parameter *γ* is the slope of distortion, with *γ* < 1 corresponding to inverted-*S*-shaped distortion, *γ*= 1 to no distortion, and *γ*>1 to *S*- shaped distortion. The parameter *p*_0_ is the crossover point where *π*(*p*)= *p*.

For each subject and condition, we fit the reported relative-frequency to the LLO model (Eq. 2) and used the estimated 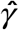 and 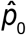 as the measures of relative-frequency distortions.

#### Response time

The response time (RT) was defined as the interval between the response screen onset and subjects’ first mouse move. The RT of the first trial of three subjects was mis-recorded due to technical issues and was excluded from further analysis. We divided all trials evenly into 5 bins based on the to-be-reported *p*, and computed the mean *p* and mean RT for each bin. A 5 (*p* bins) by 2 (numerosity conditions) repeated-measures ANOVA was performed on the mean RTs. A similar binning procedure was applied to *p*(1–*p*) to visualize the relationship of RT to *p*(1–*p*).

#### Non-parametric measures of probability distortion

As a complement to the LLO model (Eq. 2), we used a non-parametric method to visualize the probability distortion curve for each subject. In particular, we smoothed *π*(*p*)-*p* using a kernel regression method with the commonly-used Nadaraya-Watson kernel estimator [52–54]:

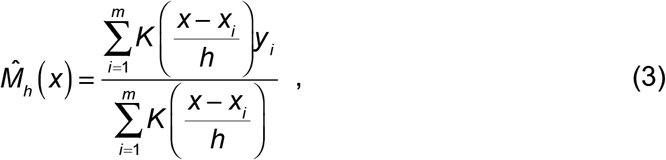

where *x_i_* and *y_i_* (*i* =1,2,...,*m*) denote observed pairs of stimuli and responses, 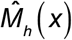 denotes the smoothed response at the stimulus value *x*, and *h* is a parameter that controls the degree of smoothing and were set to be 0.03. The *K*(·) denotes the Gaussian kernel function:

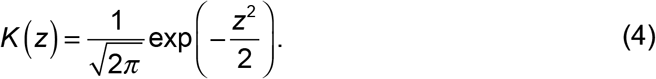

#### Model fitting

For each subject, we fit the BLO model to the subject’s *π*(*p*) in each trial using maximum likelihood estimates. The *fminsearchbnd* (J. D’Errico), a function based on *fminsearch* in MATLAB (MathWorks), was used to search for the parameters that minimized negative log likelihood.

We compared the goodness-of-fit of the BLO model with all of the alternative models based on the Akaike information criterion with a correction for sample sizes (AICc) [20, 21] and group-level Bayesian model selection [22–24]. To verify that we had found the global minimum, we repeated the searching process for 1000 times with different starting points.

### MEG Acquisition and Preprocessing

Subjects’ brain activity was recorded by a 306-channel whole-head MEG system (Elekta-Neuromag, 102 magnetometers and 102 pairs of orthogonal planar gradiometers). Head position was measured before each block by an isotrack polhemus system with four head position indicator coils (two on the left and right mastoid, the other two on the left and right forehead below the hairline). Subjects whose between-block head movement exceeded 3 mm would be excluded from further analysis. Horizontal and vertical Electro-oculograms were recorded to monitor eyemovement artifacts. Sampling rate was set to be 1000 Hz and an analog band-pass filter from 0.1 to 330 Hz was applied. Maxwell filtering was used to minimize external magnetic interference and to compensate for head movements [55, 56].

Standard preprocessing procedures were applied using Matlab 2016b and the FieldTrip package [57]. The MEG data of each block was first low-pass filtered below 20-Hz and then segmented into epochs of 7.6 s relative to trial onset (–0.6 s to 7 s). Independent component analysis (ICA) was applied to aggregated epoch data to remove artifacts including blinks, eye movements, breaths and heart activity. No subject was excluded for excessive head movement or other artifacts. For two subjects, the first trial of one block was excluded due to the failure of synchronization between stimulus onset and MEG recording onset at the beginning of the block.

### Phase Coherence Analysis

Given a periodically-changed stimulus sequence (*p* in *P*-cycle or *p*(1–*p*) in *U*-cycle), stimulus-evoked brain responses imply that the phase difference between the stimulus and response time series should be coherent across trials at the stimulus frequency [58]. The phase coherence across trials is defined as [26]:

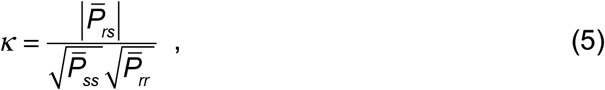

where 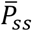 and 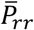 respectively denote the trial-averaged power spectrum for stimulus and response time series, and 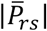 denotes the magnitude of the trial-averaged cross-spectrum between stimuli and responses. The value of *κ* for any specific frequency is between 0 and 1, with larger *κ* indicating stronger phase-locked responses.

For a specific variable (*p* or *p*(1–*p*)), we computed the phase coherence between stimulus sequence and MEG signals separately for each subject, each magnetometer and each cycle and numerosity condition. In particular, we down-sampled epoched MEG time series to 300 Hz, applied zero padding and Hanning window to single trials, and used Fast Fourier Transformation (FFT) to calculate power spectrum. The resulting frequency resolution was 0.03 Hz.

To evaluate the chance-level phase coherence, we shuffled the MEG time series across different time points within each trial and computed phase coherence for the permutated time series. This permutation procedure was repeated for 500 times to produce a distribution of chance-level phase coherences.

### Decoding Analysis

#### Time-resolved decoding

For an aperiodic stimulus sequence (*p*(1–*p*) in *P*-cycle or *p* in *U*-cycle), we could infer the brain’s encoding of the stimulus variable at a specific time lag to stimulus onset by examining how well the value of the variable could be reconstructed from the neural responses at the time lag. In particular, we performed a time-resolved decoding analysis using regression methods [28, 29, 59, 60]:

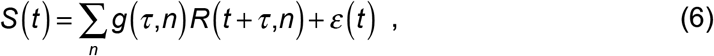

where *S*(*t*) denotes the value of the stimulus that starts at time *t*, *g*(*τ,n*) denotes the to-be-estimated decoding weight for channel *n* at time lag, *R*(*t + τ,n*) denotes the response of channel *n* at time *t*+*τ*, and *ε*(*t*) is a Gaussian noise term.

We used the MEG signals of all 306 sensors (102 magnetometers and 204 gradiometers) at single time lags to decode *p* from *U*-cycle trials or *p*(1–*p*) from *P*-cycle trials, separately for each subject, each cycle and numerosity condition, and each time lag between 0 and 900 ms. Epoched MEG time series were first down-sampled to 120 Hz. To eliminate multicollinearity and reduce overfitting, we submitted the time series of the 306 sensors to a principal component analysis (PCA) and used the first 30 components (explaining approximately 95% variance) as the regressors. The decoding weights (i.e. regression coefficients) were estimated for normalized stimuli and responses using the L2-rectified regression method implemented in the mTRF toolbox [28], with the ridge parameter set to 1.

We used a leave-one-out cross-validation [61] to evaluate the predictive power of decoding performance as the following. Out of the 50 trials in question (or 49 trials in the case of trial exclusion), each time one trial served as the test set and the remaining trials as the training set. The decoding weights estimated from the training set were applied to the test set, for which Pearson’s *r* was calculated between the predicted and the ground-truth stimulus sequence. Such computation was repeated for each trial as the test set and the averaged Pearson’s *r* was used as the measure for decoding performance.

We identified time windows that had above-chance decoding performance using cluster-based permutation tests [62] as follows. Adjacent time lags with significantly positive decoding performance at the uncorrected significance level of .05 by right-sided one-sample *t* tests were grouped into clusters and the summed *t*-value across the time lags in a cluster was defined as the cluster-level statistic. We randomly shuffled the stimulus sequence of each trial, performed the time-resolved decoding analysis on the shuffled sequence and recorded the maximum cluster-level statistic. This procedure was repeated for 500 times to produce a reference distribution of chance-level maximum cluster-level statistic, based on which we calculated the *P* value for each cluster in real data. This test effectively controls the Type I error rate in situations involving multiple comparisons. For each subject, we defined the median of all time points for which the decoding performance exceeded the 95% percentile of the distribution as the time lag of the peak [63].

#### Spatial decoding

For the conditions and time windows that had above-chance performances in the time-resolved decoding analysis, we further performed a spatial decoding analysis at individual sensor location to locate the brain regions that are involved in encoding the variable in question. For a specific senor location, the MEG signals at its three sensors (one magnetometer and two gradiometers) within the time window were used for decoding. The time window that enclosed the peak in the time-resolved decoding was 125-ms and 167-ms wide respectively for *p* and *p*(1-*p*) under *N*-large condition. The decoding procedure was similar to that of the time-resolved decoding described above, except that the number of PCA component used was 19 and 24 respectively for decoding *p* and *p(1–p)*, which explained approximately 99% variance of the original 48 (16 time lags × 3 sensors) and 63 (21 time lags × 3 sensors) channels.

### Statistical Analysis

For phase coherence or decoding performance, grand averages across subjects were computed and permutation tests described above were used to assess their statistical significance over chance [62]. The false discovery rate (*FDR*) method [64, 65] was used for multiple comparison corrections whenever applicable. For the phase coherence spectrum averaged across magnetometers, *FDR* corrections were performed among the 235 frequency bins within 0–7 Hz. For the phase coherence topography at 3.33 Hz, *FDR* corrections were performed among all 102 magnetometers.

## Supporting Information

**S1 Text. Bounded Log-Odds (BLO) model.**

**S2 Text. Factorial model comparison.**

**S3 Text. Autocorrelation analysis for periodic and aperiodic stimulus sequences.**

**S4 Text. Phase coherence and time-resolved decoding analyses for confounding factors.**

**S5 Text. Explaining the “bumps” in the phase coherence spectra of the confounding factors.**

**S6 Text. Explaining the higher phase coherence and decoding performance for *p* or *p* (1 – *p*) in the *N*-large condition than in the *N*-small condition.**

**S7 Text. Time-resolved decoding analyses based on cross-validated version of confound regression (CVCR).**

**S1 Fig. Individual subjects’** π(p) − *p* **as a function of *p*.**

The deviation of the subjective from objective relative-frequency, *π*(*p*) − *p*, is plotted as a function of *p*, separately for the *N*-small (left panel) and *N*-large (right panel) conditions of each subject. Black curves denote smoothed data. Red and blue curves respectively denote BLO and LLO model fits.

**S2 Fig. Results of model recovery analysis.**

Model parameters had been estimated for individual subjects and the 22 subjects’ fitted models were used to generate synthetic datasets of 22 virtual subjects. We generated 50 synthetic datasets for each of the 18 models considered in our factorial model comparison analysis. **(A) Model identifiability analysis.** We fit all the 18 models to each synthetic dataset and identified the best fitting model. The heatmaps represent confusion matrix that quantifies how likely a specific model was correctly identified as the best fitting model for the datasets generated by itself and how likely mistakenly as the best model for the datasets generated by the other models. Each column is for one specific model that was used to generate the datasets. Each row is for one model that was fit to the datasets generated by 18 models. **Top:** Summed ΔAICc was used to identify the best model for each dataset. The color of each cell codes the proportion that the model on its row was identified as the best model for the 50 datasets generated by the model on its column. Synthetic datasets that were generated by BLO were all best fit by BLO (see the leftmost column). Out of the 850 datasets generated from the other 17 models, only 0.24% were mis-identified to BLO (see the bottom row). **Bottom**: The group-level Bayesian model selection was used to quantify the probability for each specific model to be the best model for the dataset. The color of each cell codes the mean protected exceedance probability of the model on its row had for the 50 datasets generated by the model on its column. Higher value is coded as more reddish and lower value as more bluish. Values in each column add up to 1. From left (bottom) to right (top), the 18 models are 111, 112, 113, 121, 122, 123, 211, 212, 213, 221, 222, 223, 311, 312, 313, 321, 322, 323, where the first digit indexes the D1 assumption (1 for log-odds, 2 for Prelec, and 3 for linear), the second digit indexes the D2 assumption (1 for bounded and 2 for bounds-free), and the third digit indexes the D3 assumption (1 for 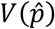 with *n*) = *b* + *N^a^*, 2 for 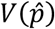 with constant *n_s_*, and 3 for constant 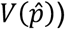. The BLO model with sample size *n_s_* = *b* + *N^a^* is the first model (111), corresponding to the leftmost column and the bottom row, which is indicated by an arrow in the plot. We can see the synthetic data that were generated by BLO were all best by BLO (see leftmost column) and those generated by the other models were seldom best fit by BLO (see bottom row). **(B) Parameter recovery analysis for BLO.** For the 50×22=1100 virtual subjects generated by the BLO model with *n_s_* = *b* + *N^a^*, the recovered parameters of BLO are plotted against the estimated parameters that were used to generate synthetic data. Each panel is for one parameter. Each dot is for one virtual subject. The parameters *η, α* and *b* that had positively skewed distributions were transformed into log scale for better visualization. A small proportion of extreme values (1.55%, 1.55% and 0.19%, respectively for *η, α* and *b*) are outside the display range. The value of *r_s_* on each panel indicates Spearman’s correlation coefficient between the estimated and recovered parameters.

**S3 Fig. Estimated parameters for the BLO model.**

In the box plot, the middle line denotes the median estimate across subjects, the bottom and top lines respectively denote the lower and upper quartiles, and the error bars denote the 99% confidence interval. Dots denote estimates for individual subjects. The parameters *η, α* and *b* were transformed into log scale for better visualization.

**S4 Fig. Autocorrelations of stimulus sequences as functions of time lags.**

Left: the *P*-cycle condition. Right: the *U*-cycle condition. Solid and dashed lines are respectively for the *p* and *p*(1-*p*) sequences. The stimulus sequences used for autocorrelation calculations were from the representative subject whose behavior results are showed in Fig 2A. All other subjects had similar autocorrelation patterns.

**S5 Fig. Results of control analysis for the confounding variables.**

**(A) Correlation analysis.** Top row: Pearson’s correlations between the *p* sequence and the eight confounding variables (*N*, *N_t_*, *N_o_*, *AvgLumi*, *vCIE-L\**, *vCIE-a\**, *vCIE-b\** and *M-contrast*) separately for the four cycle and numerosity conditions. Bottom row: Pearson’s correlations between the *p*(1–*p*) sequence and the eight confounding variables for each condition. The results confirmed our design that all of the confounding variables had negligible or moderate correlations with *p* and *p*(1–*p*). **(B) Phase coherence analysis.** Grand-averaged phase coherence spectrum between the sequences of the eight confounding variables and neural responses from magnetometers. Here we have reproduced the results of the phase coherence analysis for *p* and *p*(1–*p*) (inside the black frame) to facilitate a comparison between the main analysis and the control analysis. The observed phase coherence spectra of the confounding variables had different patterns than those of *p* or *p*(1–*p*). The vertical line marks 3.33 Hz. See S5 Text and S6 Fig for the possible cause of the “bumps” between 0 and 6.67 Hz. **(C) Time-resolved decoding analysis.** The same time-resolved decoding procedures as Fig 5 were applied to the eight confounding variables. Here we have reproduced the results of the time-resolved decoding analysis for *p* and *p*(1–*p*) (inside the black frame) to facilitate a comparison between the main analysis and the control analysis. The time-resolved decoding performance of the confounding variables had different patterns than those of *p* or *p*(1–*p*). For **(A)**–**(C)**, light and dark green respectively code the *N*-small and *N*-large conditions. For **(B)** and **(C)**, solid and dashed lines respectively denote the *P*-cycle and *U*-cycle conditions. Shadings denote SEM across subjects.

**S6 Fig**. **Explaining the “bumps” in the phase coherence of numerosity.**

Simulated neural signals were generated through convolution of TRF estimated from the MEG2521 sensor (located right occipital) with the *N* sequence, perturbed by Gaussian nose whose standard deviation was set to 4. The simulated neural signals were then submitted to the phase coherence analysis (see Materials and Methods). Light and dark green curves denote the phase coherence across magnetometers of real data from one example subject. Orange curves denote the average across 500 simulations. The “bump” across 0–6.67 Hz observed in real data was reproduced in the simulated data.

**S7 Fig**. **Spatial decoding topography for the other three statistically significant time windows in time-resolved decoding analysis.** Black dots mark the sensor locations with above-chance decoding performance (*P_FDR_* < 0.05).

**S8 Fig. Individual subjects’ time-resolved decoding performance for *p* (left column) and *p*(1–*p*) (right column), separately for the *N*-small (top row) and *N*- large (bottom row) conditions.**

Each row on the heatmap is for one subject. Higher level of decoding performance is coded as more reddish and lower level as more bluish. The green curve superimposed on the heatmap denotes grand-averaged decoding performance across subjects (see the right y-axis). Symbols on the top of each panel indicate time lags that had above-chance decoding performance (cluster-based permutation tests), with vertical bars representing *P_cluster_*<0.01 and dots representing 0.01≤*P_cluster_*<0.05.

**S9 Fig. Time-resolved decoding analysis based on cross-validated version of confound regression (CVCR).**

**(A)** Decoding performance for *p*(1-*p*). **(B)** Decoding performance for 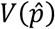. The 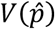 *was* computed according to each subject’s fitted BLO model (with sample size *n_s_* = *b* + *N^α^*). Curves denote grand-averaged decoding performance over different time lags, separately for the *N*-small (light green) and *N*-large (dark green) conditions. Shadings denote SEM across subjects. Vertical bars above the curves indicate time lags that had above-chance decoding performance (cluster-based permutation tests, *P_cluster_*<0.01). The eight confounding variables we considered in S4 Text (*N*, *N_t_*, *N_o_*, *AvgLumi*, *vCIE-L\**, *vCIE-a\**, *vCIE-b\** and *M-contrast*) were regressed out from MEG time series before decoding analysis. See S7 Text for methodological details. The decoded time course from CVCR for *p*(1–*p*) was similar to that of the standard time-resolved decoding analysis (Fig 5, right panel). The decoded time course was also similar for 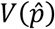.

**S10 Fig. Consequences of random sampling errors.**

The *p* or *p*(1–*p*) estimated from a sample of dots from a display might deviate from the true value of *p* or *p*(1–*p*) in the display. As an evaluation of the fidelity of the sampled value, we used numerical simulations to compute the correlation (Pearson’s *r*) between the true value and the sampled value, based on the BLO parameters estimated from individual subjects. Triangles and circles respectively denote *p* and *p*(1–*p*). The truth-sample correlation was higher in the *N*-large condition than in the *N*-small condition.

