## Supporting Information for "Automatic and fast encoding of representational uncertainty underlies probability distortion"

### S1 Text. Bounded Log-Odds (BLO) model

BLO is based on the following three assumptions.

**1. Probability is internally represented in log-odds.** BLO assumes that any probability or relative frequency, *p*, is internally represented as a linear transformation of log-odds,

$$\lambda(p) = \log\frac{p}{1-p} \ , \tag{S1}$$

which potentially extends from minus infinity ( $\lambda(0) = -\infty$ ) to infinity ( $\lambda(1) = \infty$ ).

**2. Representation is encoded on a bounded Thurstone scale.** By "Thurstone scale", we refer to a psychological scale perturbed by independent, identically-distributed Gaussian noise, which was proposed by Thurstone [1] and has been widely used in modeling representations of psychological magnitudes [e.g., 2, 3]. BLO assumes that the Thurstone scale for encoding log-odds has a limited range $[-\Psi, \Psi]$ and can only encode a selected interval of log-odds. It defines bounding operation

$$\Gamma[\lambda] = \begin{cases} \Delta^-, & \lambda < \Delta^- \\ \lambda, & \Delta^- \le \lambda \le \Delta^+ \\ \Delta^+, & \lambda > \Delta^+ \end{cases} \tag{S2}$$

to confine the representation of log-odds $\lambda$ to the interval $[\Delta^-, \Delta^+]$, where $\Delta^-$ and $\Delta^+$ are free parameters. That is, any $\lambda(p)$ that is outside the bounds will be truncated to its nearest bound. The bounded interval $[\Delta^-, \Delta^+]$ is mapped to the bounded Thurstone scale $[-\Psi, \Psi]$ through a linear transformation, so that the log-odds $\lambda(p)$ is encoded as

$$\Lambda(p) = \eta\left[\Gamma(\lambda[p]) - (\Delta^- + \Delta^+)/2\right], \tag{S3}$$

where $\eta \equiv \dfrac{\Psi}{(\Delta^+ - \Delta^-)/2}$ is a free parameter.

**3. Representational uncertainty is compensated in the final estimate of probability.** In our relative-frequency judgment task, we assume representational uncertainty partly arises from random variation of sampling. Suppose subjects may not access to all of the dots in a display. For a display of $N$ dots, a sample of $n_s$ dots is randomly drawn from the display without replacement and used to infer the relative frequency. The resulting variance of $\hat{p}$ is thus [4, 5]:

$$V(\hat{p}) = \frac{p(1-p)}{n_s}\frac{N-n_s}{N-1} . \tag{S4}$$

In the present study, $n_s$ is modeled as an increasing function of numerosity:

$$n_s = b + N^a , \tag{S5}$$

where $a \geq 0$ and $b \geq 0$ are free parameters. To keep the resulting $V(\hat{p}) \geq 0$, we forced $V(\hat{p}) = 0$ whenever $V(\hat{p}) < 0$. That is, $n_s = b + N^a$ is the maximum number of dots one can integrate in one's judgment. Note that Eq. S4 does not necessarily imply that $V(\hat{p})$ is an increasing function of $N$, given that $n_s$ is not constant but may vary with $N$ (Eq. S5). In fact, for a median subject ($a$=0.42 and $b$=1.91) in our experiment, $V(\hat{p})$ may slightly increase with $N$ for very small $N$ (Fig 2C, from $N$=10 to $N$=20) but mostly be a *decreasing* function of $N$ (Fig 2C, from $N$=20 to $N$=900).

BLO assumes that representational uncertainty is compensated in the final estimate of probability, with the log-odds of $\pi(p)$ modeled as a weighted average of $\Lambda(p)$ and a fixed point $\Lambda_0$ on the log-odds scale:

$$\lambda\left[\pi(p)\right] = \omega_p \Lambda(p) + \left(1-\omega_p\right)\Lambda_0 + \varepsilon_\lambda , \tag{S6}$$

where $\omega_p = \frac{1}{1+\alpha V(\hat{p})}$ is a measure of the reliability of the internal representation, and $\varepsilon_\lambda$ is Gaussian noise term on the log-odds scale with mean 0 and variance $\sigma_\lambda^2$. Here $\alpha \geq 0$ and

$\sigma_\lambda > 0$ are free parameters. In total, the BLO model for our relative-frequency judgment task has eight parameters: $\Delta^-$, $\Delta^+$, $a$, $b$, $\eta$, $\Lambda_0$, $\alpha$, and $\sigma_\lambda$.

#### S2 Text. Factorial model comparison

Similar to Zhang et al. [1], we performed a factorial model comparison [2] to test whether each of the three assumptions in the BLO model outperformed plausible alternative assumptions in fitting behavioral data. The models we considered differ in the following three dimensions.

D1: scale of transformation. The scale on which probability is internally represented can be the log-odds scale ($\lambda(p) = \log\frac{p}{1-p}$ as in S1 Text, Eq. S1), the Prelec scale ($\lambda'(p) = -\log(-\log p)$) derived from the two-parameter Prelec function [3], or the linear scale used in the neo-additive probability distortion functions [see 4 for a review]. See Zhang et al. [1] for more details.

D2: bounded versus bounds-free, concerning whether the bounding operation (S1 Text, Eq. S2) is involved.

D3: variance compensation. BLO compensates for representational uncertainty following S1 Text, Eq. S6. In relative-frequency judgment, the form of $V(\hat{p})$ is not only proportional to $p$(1–$p$), but also depends on $N$ and sampling strategy. In the present study, we modeled sample size as $n_s = b + N^a$, which increases with $N$. Alternatively, sample size $n_s$ may be modeled as a constant. Note that under constant $n_s$ the value of is still proportional to *p(1–p)*. A third alternative assumption is to set $V(\hat{p}) = \text{constant}$, which is effectively implemented in classic descriptive models of probability distortion such as LLO or the Prelec function.

These three dimensions are independent of each other, analogous to the different factors manipulated in a factorial experimental design. In total, there are 3 (D1: log-odds, Prelec, or linear) × 2 (D2: bounded or bounds-free) × 3 (D3: $V(\hat{p})$ with $n_s = b + N^a$, $V(\hat{p})$ with constant $n_s$, or constant $V(\hat{p})$) = 18 models.

In this three-dimensional model space, the BLO model we introduced earlier corresponds to (D1=log-odds, D2=bounded, D3=$V(\hat{p})$ with $n_s = b + N^a$). The model at (D1=log-odds, D2=bounded, D3=$V(\hat{p})$ with constant $n_s$) is also a BLO model, which differs from our main BLO model in the specification of sampling strategies. For simplicity, in presenting the results of factorial model comparison, we will only refer to our main BLO model as BLO. The LLO and two-parameter Prelec models are also special cases of the 18 models, respectively corresponding to (D1=log-odds, D2=bounds-free, D3=constant $V(\hat{p})$) and (D1=Prelec, D2=bounds-free, D3=constant $V(\hat{p})$).

#### S3 Text. Autocorrelation analysis for periodic and aperiodic stimulus sequences

We applied an autocorrelation analysis to the stimulus sequences in each trial to confirm their periodicity or aperiodicity. Autocorrelations were calculated for individual trials and then averaged across trials in each condition. As designed, the $p$ sequence in $P$-cycle trials and the $p(1–p)$ sequence in $U$-cycle trials had periodic autocorrelations, while the $p$ sequence in $U$-cycle trials and the $p(1–p)$ sequence in $P$-cycle trials were free of autocorrelations (S4 Fig).

#### S4 Text. Phase coherence and time-resolved decoding analyses for confounding factors

We considered the following eight variables in the stimuli as potential confounding factors, which may covary with *p* or *p*(1–*p*) across displays.

(1) *N*: the total number of dots (numerosity) in the display, which by design was independent of both *p* and *p*(1–*p*).

(2) $N_t$: the number of dots in the target color in the display, that is, $N_t = Np$.

(3) $N_o$: the number of dots in the other color in the display, that is, $N_o = N(1-p)$.

(4) *AvgLumi*: the mean luminance of the display, which was kept constant across different displays and different trials and was thus independent of both *p* and *p*(1–*p*).

(5) *vCIE-L**: the color variance of the display on the L* dimension in the CIELAB color space, where L* ranges from black to white.

(6) *vCIE-a**: the color variance of the display on the a* dimension in the CIELAB color space, where a* ranges from green to red.

(7) *vCIE-b**: the color variance of the display on the b* dimension in the CIELAB color space, where b* ranges from blue to yellow.

(8) *M-contrast*: the *Michelson* luminance contrast [1], that is, *M-contrast*=$\frac{L_{dots}-L_{background}}{L_{dots}+L_{background}}$.

According to our design, most of these variables had negligible correlations with *p* and *p(1–p)* and none of the absolute values of correlation coefficients exceeded 0.53 (Pearson's *r*, S5A Fig). The numerosity (*N*) had considerable correlations (Pearson's $|r| > 0.79$) with all other confounding factors except for the mean luminance (*AvgLumi*), which was constant across displays. We performed similar phase coherence (S5B Fig) and time-resolved decoding analyses (S5C Fig) for these variables as we did for *p* and *p*(1–*p*) but obtained very different patterns, which suggests that the findings we reported for *p* and *p*(1–*p*) in the main text are unlikely to be effects of confounding factors. The patterns of all these confounding factors expect for *AvgLumi* were almost identical, which may have the same origins such as the automatic encoding of numerosity [2, 3].

### S5 Text. Explaining the "bumps" in the phase coherence spectra of the confounding factors

As shown in the S5B Fig, there was a "bump" spanning across 0–6.67 Hz in the phase coherence spectrum of *N*. So did all of the confounding variables we considered above that had high correlations with *N*. Though also centering at 3.33 Hz, these "bumps" were much wider than the 3.33 Hz peak we observed for the *p* sequence in the *P*-cycle condition or the *p*(1–*p*) sequence in the *U*-cycle condition. Besides, the "bumps" were similar across all four cycle and numerosity conditions, which were unlikely due to the periodicity of *p* or *p*(1–*p*). We found in a simulation analysis that these "bumps" could simply result from the particular temporal course of the neural responses to the variables.

In particular, we first estimated the temporal course of stimulus-evoked brain responses to *N* using a computational technique known as Temporal Response Function (TRF) analysis [1-3]. The TRF for a specific stimulus dimension describes how the brain responds to a unit change in stimulus magnitude at different time lags. The brain is modeled as a linear system, whose response at a specific time point is the summation of the responses to all previous stimuli, that is, a convolution of the stimulus sequence with TRF plus Gaussian random noise:

$$R(t,n) = \sum_{\tau} w(\tau,n) S(t-\tau) + \varepsilon(t,n), \tag{S7}$$

where *t* denotes time, *n* denotes sensor number, $\tau$ denotes the time lag to stimulus onset, $R(\bullet)$, $w(\bullet)$, $S(\bullet)$ and $\varepsilon(\bullet)$ respectively denote the magnitude of sensor response, the to-be-estimated TRF, the stimulus sequence, and Gaussian noise term. The stimulus sequence was modeled as the summation of pulses at display onsets. For the TRF analysis, epoched MEG data were down-sampled to 120 Hz. The TRF $w(\tau,n)$ was estimated for a maximum delay of 900 ms using a rectified regression method, with the ridge parameter set to 1. Both the stimulus sequence and sensor responses were normalized before the analysis.

We estimated the TRF for *N* and then convolved it with the *N* sequence and added a Gaussian noise to generate virtual neural responses. We then applied the phase coherence analysis to the virtual neural responses. This simulation was repeated for 500 times, based on which we computed the mean phase coherence spectrum for the simulated data. We found that the resulting phase coherence spectrum resembled that of real data, reproducing the “bump” between 0 and 6.67 Hz (S6 Fig).

#### S6 Text. Explaining the higher phase coherence and decoding performance for *p* or *p*(1–*p*) in the *N*-large condition than in the *N*-small condition

In our experimental design, the values of *p* or *p*(1–*p*) were independent of the total number of dots in each display, *N*. Then why should we observe higher phase coherence and decoding performance for *p* or *p*(1–*p*) in the *N*-large condition than in the *N*-small condition? We conjectured that this difference arose from random sampling errors in subjects' estimation of *p* or *p*(1–*p*). As inferred from subjects' behavioral responses, the *N*-large condition was accompanied by a larger sample size, thus leading to a higher correlation between the true *p* or *p*(1–*p*) and that estimated from the sample (see S10 Fig for simulation results). Therefore, subjects' brain responses to *p* or *p*(1–*p*) would be less variable and more related to the objective *p* or *p*(1–*p*) in the *N*-large condition. Consequently, we observed a higher association (quantified by phase coherence or decoding performance) between MEG time series and the objective *p* or *p*(1–*p*) in the *N*-large condition than in the *N*-small condition.

#### S7 Text. Time-resolved decoding analyses based on cross-validated version of confound regression (CVCR)

To further rule out the possibility that the observed automatic encoding of $p(1-p)$ may be an artefact of confounding factors, we performed a time-resolved decoding analysis based on the cross-validated version of confound regression (CVCR) method proposed by Snoek, Miletić and Scholte [1], which allowed us to regress out confounding variables from MEG time series before decoding analysis.

The CVCR method consists two modules: encoding and decoding. During encoding, we fit a multiple linear regression model to MEG time series separately for each magnetometer or gradiometer and each specific time lag, with the confounding variables as regressors:

$$Y_j(\tau) = \sum_{i=1}^{8} \beta_{j,i}(\tau) C_i + \beta_{j,0} + \varepsilon_j(\tau) \ , \tag{S8}$$

where $j$ denotes sensor number, $\tau$ denotes time lag after stimulus onset (ranging from 0 to 900 ms), $Y_j(\tau)$ denotes MEG signals at sensor $j$ and time lag $\tau$, $C_i$ ($i$=1, 2, …, 8) denotes the value of the $i$-th confounding variable, among the eight confounding variables we considered in S4 Text (i.e., $N$, $N_t$, $N_o$, *AvgLumi*, *vCIE-L**, *vCIE-a**, *vCIE-b** and *M-contrast*), $\beta_{j,i}$ and $\beta_{j,0}$ are free parameters, and $\varepsilon_j(\tau)$ is a Gaussian noise term.

Then, we subtracted the explained variance of the confounding variables from MEG signals and defined the residual as "confounds-regressed-out MEG signal":

$$Y_{j,reg}(\tau) = Y_j(\tau) - \left( \sum_{i=1}^{8} \hat{\beta}_{j,i}(\tau) C_i + \hat{\beta}_{j,0} \right) . \tag{S9}$$

The confounds-regressed-out MEG signal, $Y_{j,reg}$, was subsequently used for time-resolved decoding analysis.

Following suggestions by Snoek, Miletić and Scholte [1], we used a leave-one-out procedure as follows. Out of the 50 trials in question (or 49 trials in the case of trial exclusion), each time one trial served as the test set and the remaining trials as the training set. The parameters (of Eq. S8) estimated within each fold of training data, $\hat{\beta}_{j,i}^{train}$ and $\hat{\beta}_{j,0}^{train}$, were used to remove the variance related to the confounds from both the training set and test set. The resulting confounds-regressed-out training data, $Y_{j,reg}^{train}$, and test data, $Y_{j,reg}^{test}$, were then concatenated and used for the decoding module. When all eight confounding variables were regressed out, the decoding performance of $p(1–p)$ was little influenced (S9A Fig). We also applied the same CVCR procedure to representational uncertainty, $V\left(\hat{p}\right)$, defined in S1 Text. The resulting decoding performance of $V\left(\hat{p}\right)$ was similar to that of $p(1–p)$ (S9B Fig).

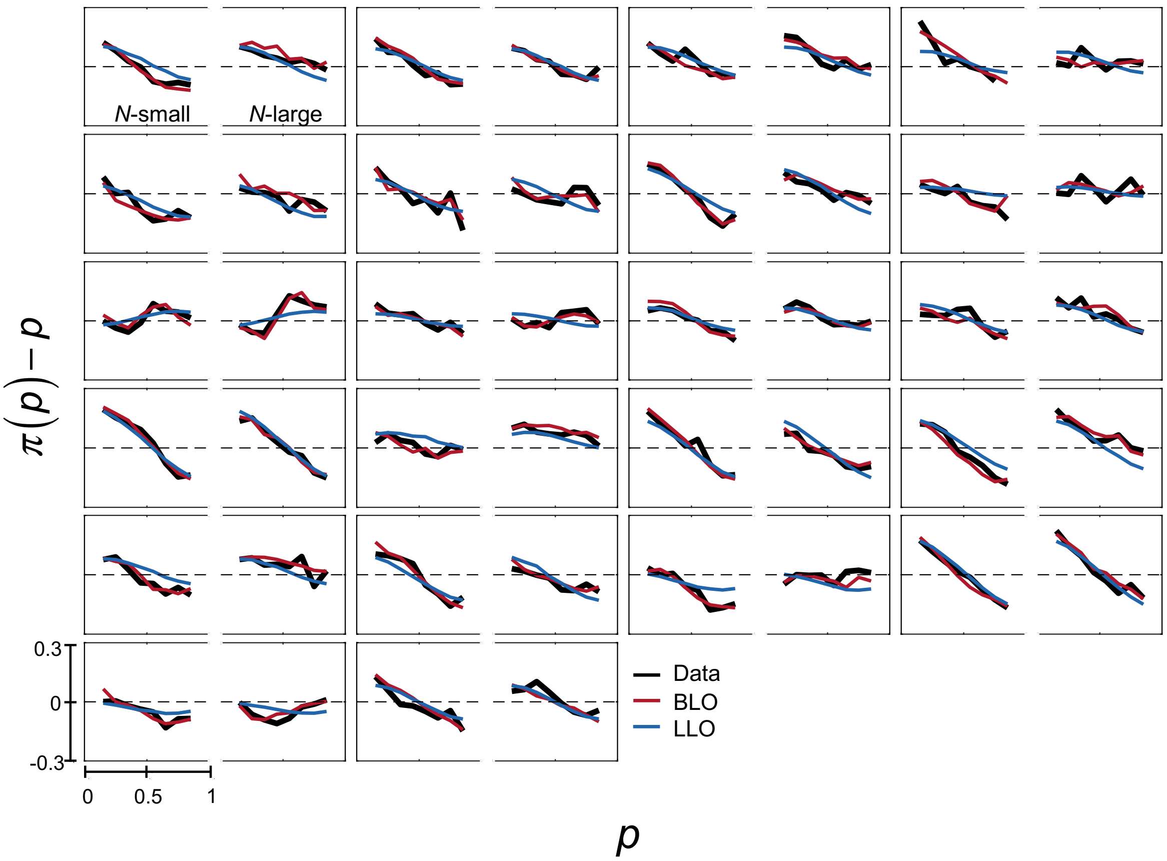


**S1 Fig. Individual subjects' $\pi(p) - p$ as a function of *p*.**

The deviation of the subjective from objective relative-frequency, $\pi(p) - p$, is plotted as a function of *p*, separately for the *N*-small (left panel) and *N*-large (right panel) conditions of each subject. Black curves denote smoothed data. Red and blue curves respectively denote BLO and LLO model fits.

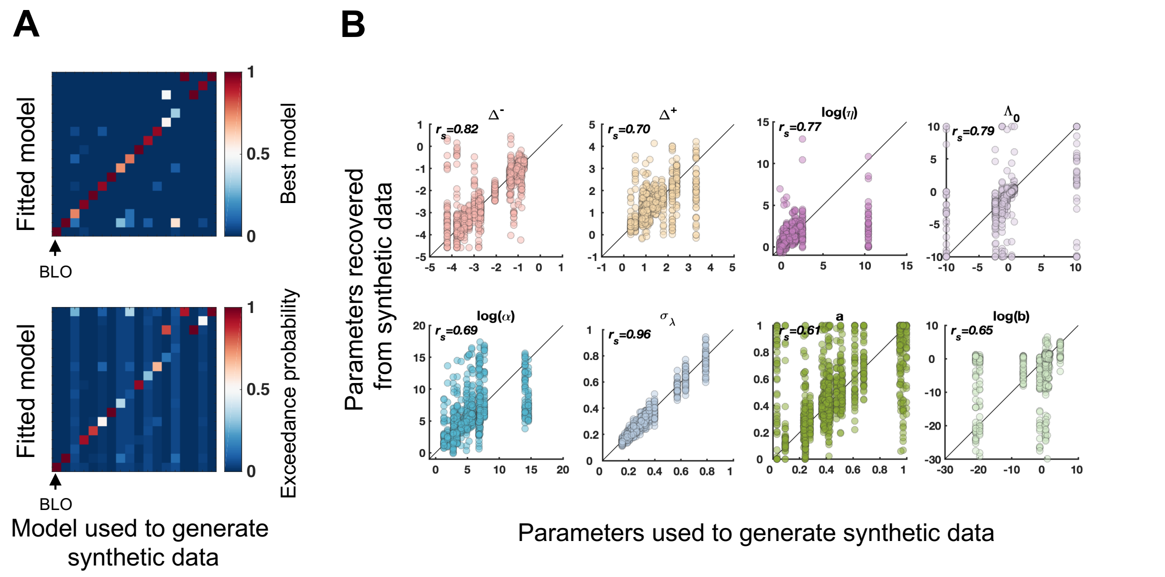


**S2 Fig. Results of model recovery analysis.**

Model parameters had been estimated for individual subjects and the 22 subjects' fitted models were used to generate synthetic datasets of 22 virtual subjects. We generated 50 synthetic datasets for each of the 18 models considered in our factorial model comparison analysis. **(A) Model identifiability analysis.** We fit all the 18 models to each synthetic dataset and identified the best fitting model. The heatmaps represent confusion matrix that quantifies how likely a specific model was correctly identified as the best fitting model for the datasets generated by itself and how likely mistakenly as the best model for the datasets generated by the other models. Each column is for one specific model that was used to generate the datasets. Each row is for one model that was fit to the datasets generated by 18 models. **Top:** Summed ΔAICc was used to identify the best model for each dataset. The color of each cell codes the proportion that the model on its row was identified as the best model for the 50 datasets generated by the model on its column. Synthetic datasets that were generated by BLO were all best fit by BLO (see the leftmost column). Out of the 850 datasets generated from the other 17 models, only 0.24% were mis-identified to BLO (see the bottom row). **Bottom**: The group-level Bayesian model selection was used to quantify the probability for each specific model to be the best model for the dataset. The color of each cell codes the mean protected exceedance probability of the model on its row had for the 50 datasets generated by the model on its column. Higher value is coded as more reddish and lower value as more bluish. Values in each column add up to 1. From left (bottom) to right (top), the 18 models are 111, 112, 113, 121, 122, 123, 211, 212, 213, 221, 222, 223, 311, 312, 313, 321, 322, 323, where the first digit indexes the D1 assumption (1 for log-odds, 2 for Prelec, and 3 for linear), the second digit indexes the D2 assumption (1 for bounded and 2 for bounds-free), and the third digit indexes the D3 assumption (1 for $V(\hat{p})$ with $n_s = b + N^a$, 2 for $V(\hat{p})$ with constant $n_s$, and 3 for constant $V(\hat{p})$). The BLO model with sample size $n_s = b + N^a$ is the first model (111), corresponding to the leftmost column and the bottom row, which is indicated by an arrow in the plot. We can see the synthetic data that were generated by BLO were all best by BLO (see leftmost column) and those generated by the other models were seldom best fit by BLO (see bottom row). **(B) Parameter recovery analysis for BLO.** For the 50×22=1100 virtual subjects generated by the BLO model with $n_s = b + N^a$ , the recovered parameters of BLO are plotted against the estimated parameters that were used to generate synthetic data. Each panel is for one parameter. Each dot is for one virtual subject. The parameters $\eta$, $\alpha$ and $b$ that had positively skewed distributions were transformed into log scale for better visualization. A small proportion of extreme values (1.55%, 1.55% and 0.19%, respectively for $\eta$, $\alpha$ and $b$) are outside the display range. The value of $r_s$ on each panel indicates Spearman's correlation coefficient between the estimated and recovered parameters.

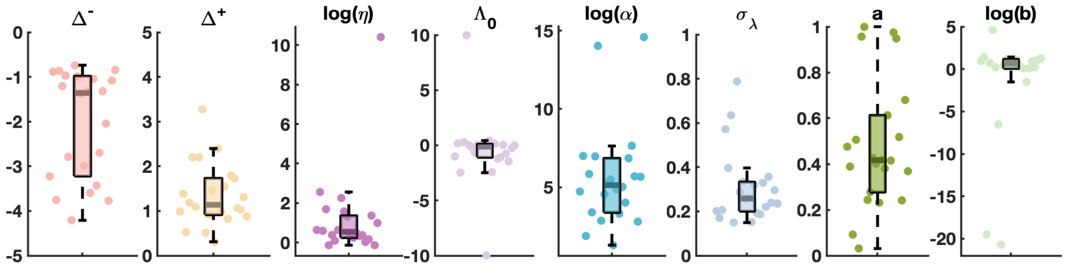


**S3 Fig. Estimated parameters for the BLO model.**

In the box plot, the middle line denotes the median estimate across subjects, the bottom and top lines denote the lower and upper quartiles, and the error bars denote the 99% confidence interval. Dots denote estimates for individual subjects. The parameters $\eta$, *a* and $b$ were transformed into log scale for better visualization.

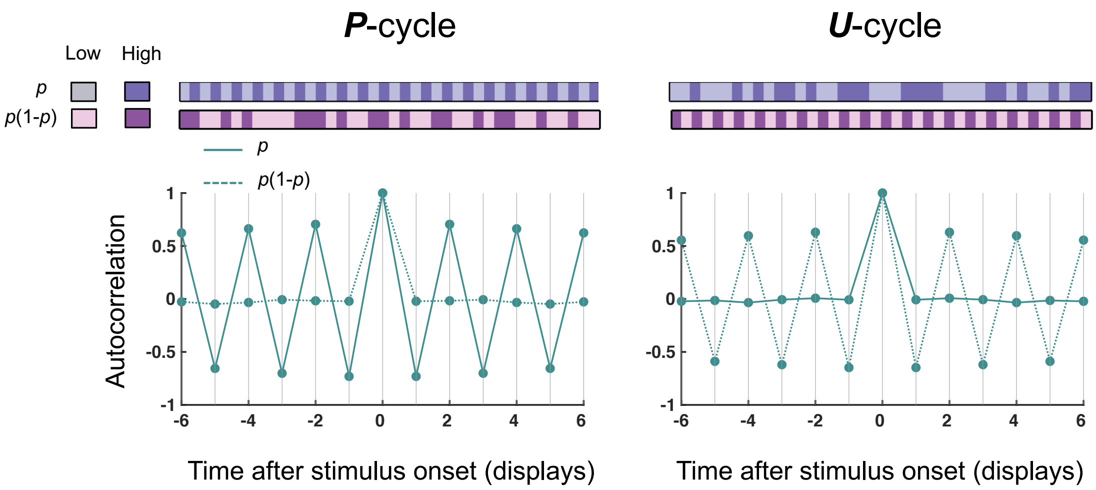


**S4 Fig. Autocorrelations of stimulus sequences as functions of time lags.**
Left: the *P*-cycle condition. Right: the *U*-cycle condition. Solid and dashed lines are respectively for the $p$ and $p(1-p)$ sequences. The stimulus sequences used for autocorrelation calculations were from the representative subject whose behavior results are showed in Fig 2A. All other subjects had similar autocorrelation patterns.

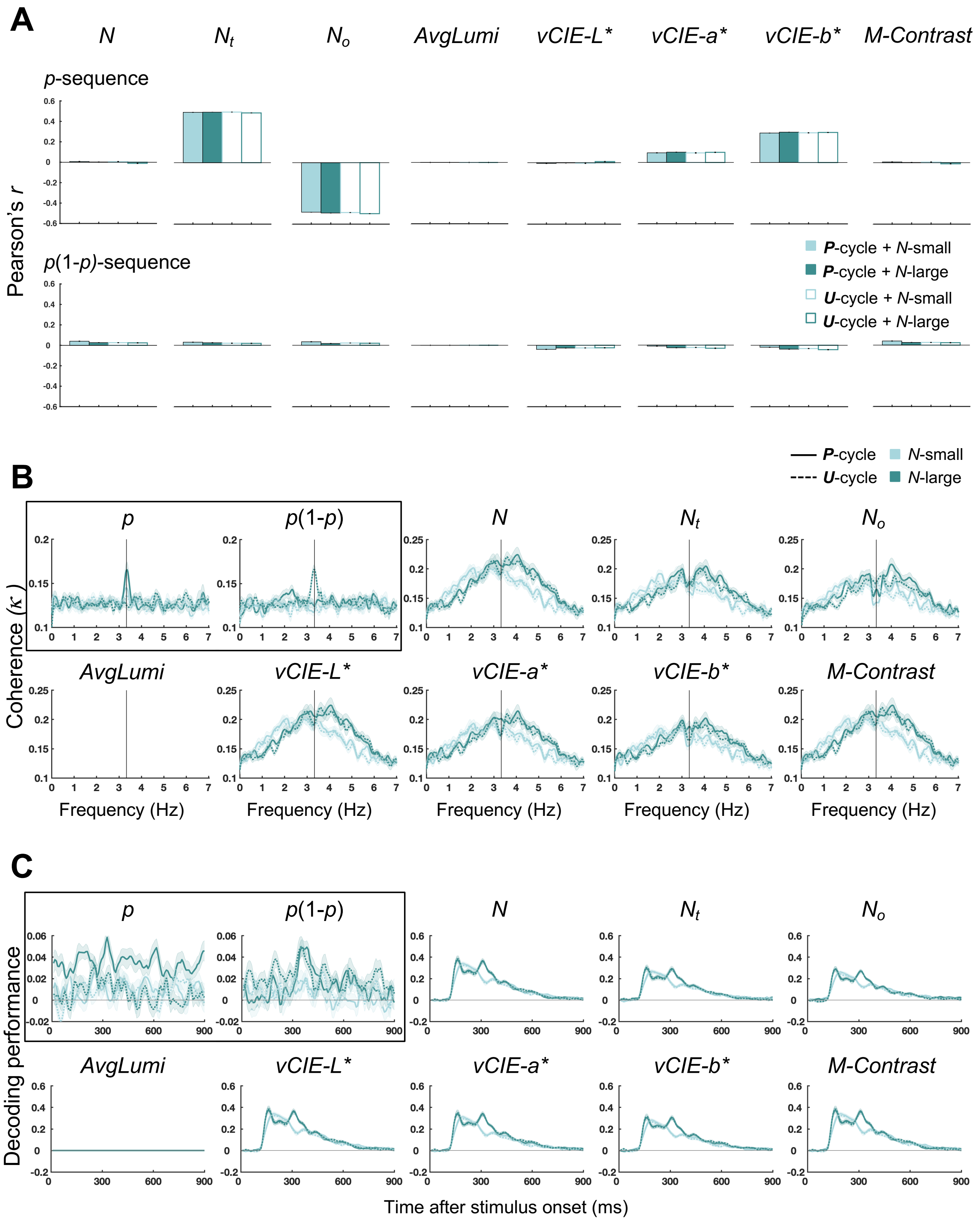


**S5 Fig. Results of control analysis for the confounding variables.**

**(A) Correlation analysis.** Top row: Pearson's correlations between the *p* sequence and the eight confounding variables (*N*, $N_t$, $N_o$, *AvgLumi*, *vCIE-L**, *vCIE-a**, *vCIE-b** and *M-contrast*) separately for the four cycle and numerosity conditions. Bottom row: Pearson's correlations between the *p*(1–*p*) sequence and the eight confounding variables for each condition. The results confirmed our design that all of the confounding variables had negligible or moderate correlations with *p* and *p(1–p)*. **(B) Phase coherence analysis.** Grand-averaged phase coherence spectrum between the sequences of the eight confounding variables and neural responses from magnetometers. Here we have reproduced the results of the phase coherence analysis for *p* and *p*(1–*p*) (inside the black frame) to facilitate a comparison between the main analysis and the control analysis. The observed phase coherence spectra of the confounding variables had different patterns than those of *p* or *p*(1–*p*). The vertical line marks 3.33 Hz. See S5 Text and S6 Fig for the possible cause of the "bumps" between 0 and 6.67 Hz. **(C) Time-resolved decoding analysis.** The same time-resolved decoding procedures as Fig 5 were applied to the eight confounding variables. Here we have reproduced the results of the time-resolved decoding analysis for *p* and *p*(1–*p*) (inside the black frame) to facilitate a comparison between the main analysis and the control analysis. The time-resolved decoding performance of the confounding variables had different patterns than those of *p* or *p*(1–*p*). For **(A)**–**(C)**, light and dark green respectively code the *N*-small and *N*-large conditions. For **(B)** and **(C)**, solid and dashed lines respectively denote the *P*-cycle and *U*-cycle conditions. Shadings denote SEM across subjects.

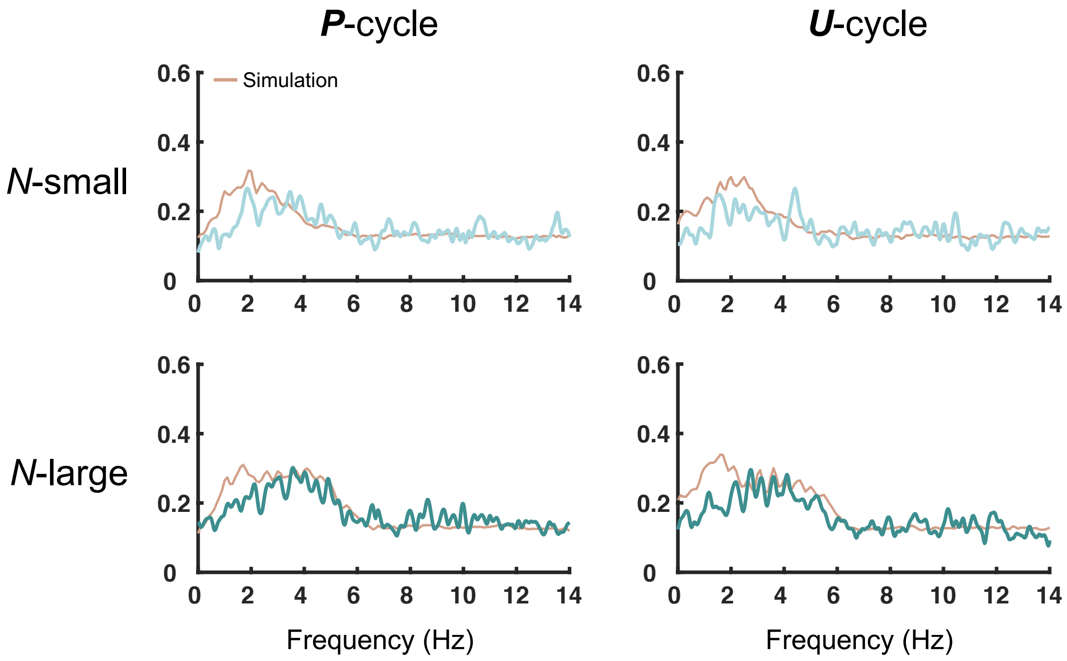


**S6 Fig. Explaining the "bumps" in the phase coherence of numerosity.**
Simulated neural signals were generated through convolution of TRF estimated from the MEG2521 sensor (located right occipital) with the *N* sequence, perturbed by Gaussian nose whose standard deviation was set to 4. The simulated neural signals were then submitted to the phase coherence analysis (see Materials and Methods). Light and dark green curves denote the phase coherence across magnetometers of real data from one example subject. Orange curves denote the average across 500 simulations. The "bump" across 0–6.67 Hz observed in real data was reproduced in the simulated data.

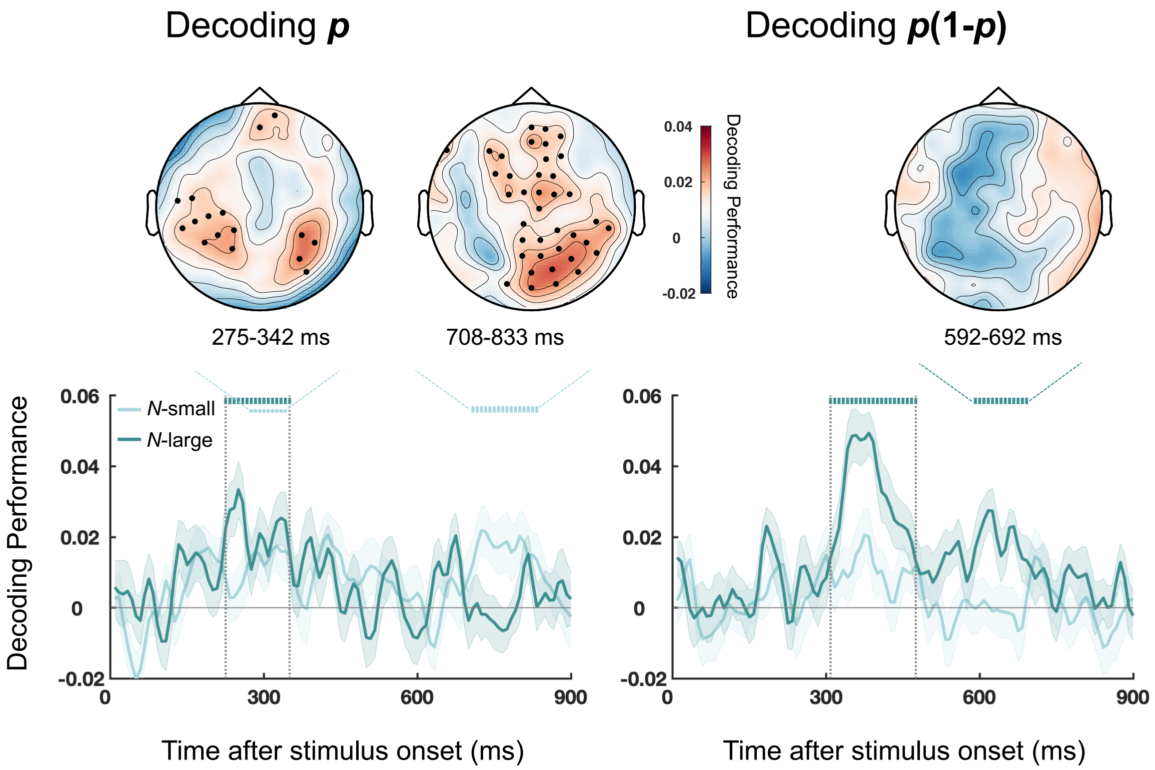


**S7 Fig. Spatial decoding topography for the other three statistically significant time windows in time-resolved decoding analysis.** Black dots mark the sensor locations with above-chance decoding performance ($P_{FDR} < 0.05$).

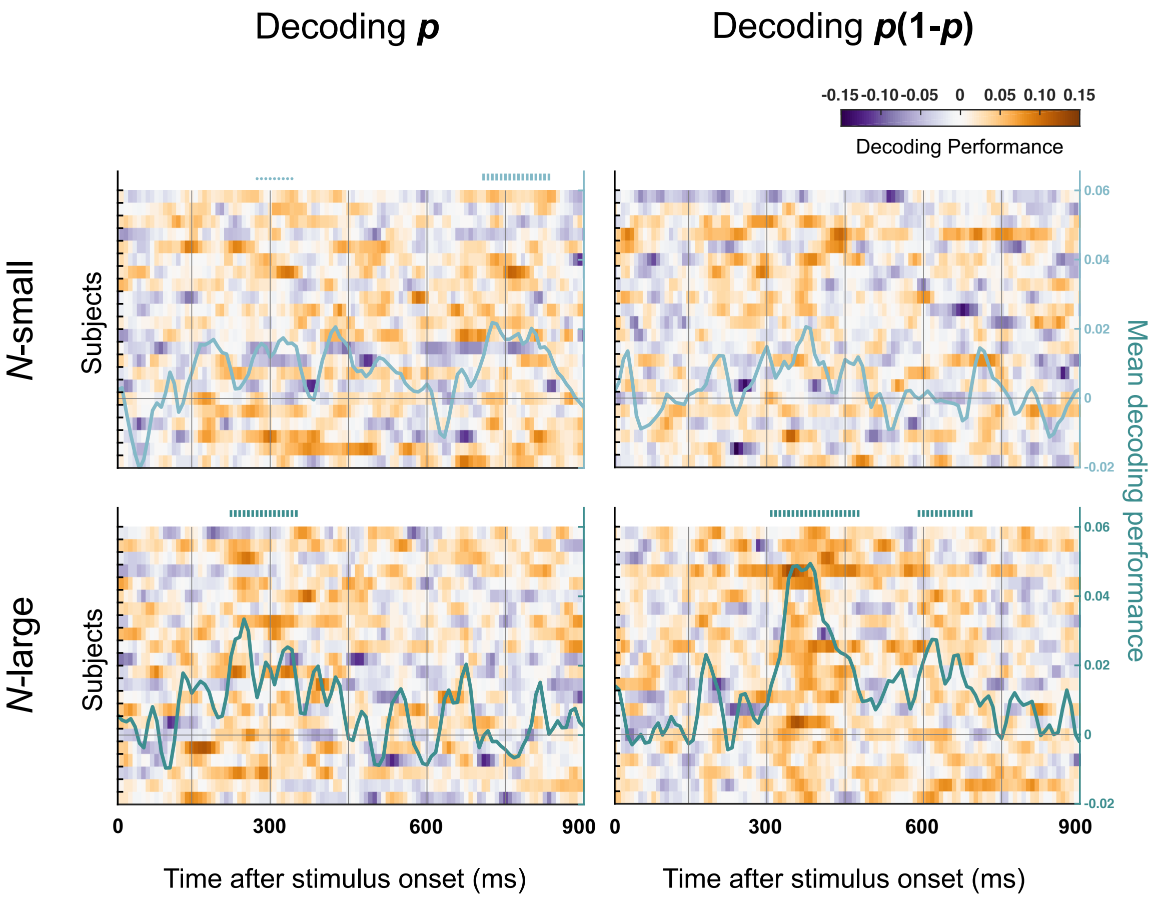


**S8 Fig. Individual subjects' time-resolved decoding performance for *p* (left column) and *p*(1–*p*) (right column), separately for the *N*-small (top row) and *N*-large (bottom row) conditions.**

Each row on the heatmap is for one subject. Higher level of decoding performance is coded as more reddish and lower level as more bluish. The green curve superimposed on the heatmap denotes grand-averaged decoding performance across subjects (see the right y-axis). Symbols on the top of each panel indicate time lags that had above-chance decoding performance (cluster-based permutation tests), with vertical bars representing $P_{cluster}<0.01$ and dots representing $0.01\leq P_{cluster}<0.05$.

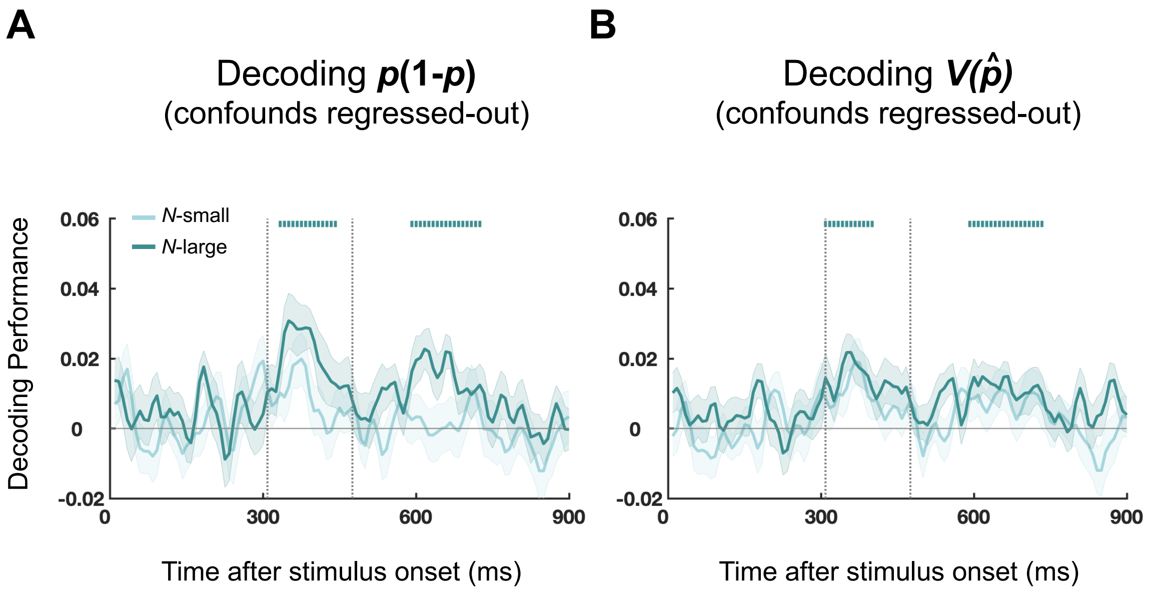


**S9 Fig. Time-resolved decoding analysis based on cross-validated version of confound regression (CVCR).**

**(A)** Decoding performance for *p*(1–*p*). **(B)** Decoding performance for $V(\hat{p})$. The $V(\hat{p})$ was computed according to each subject's fitted BLO model (with sample size $n_s = b + N^a$). Curves denote grand-averaged decoding performance over different time lags, separately for the *N*-small (light green) and *N*-large (dark green) conditions. Shadings denote SEM across subjects. Vertical bars above the curves indicate time lags that had above-chance decoding performance (cluster-based permutation tests, $P_{cluster}$<0.01). The eight confounding variables we considered in S4 Text (*N*, $N_t$, $N_o$, *AvgLumi*, *vCIE-L**, *vCIE-a**, *vCIE-b** and *M-contrast*) were regressed out from MEG time series before decoding analysis. See S7 Text for methodological details. The decoded time course from CVCR for *p*(1–*p*) was similar to that of the standard time-resolved decoding analysis (Fig 5, right panel). The decoded time course was also similar for $V(\hat{p})$.

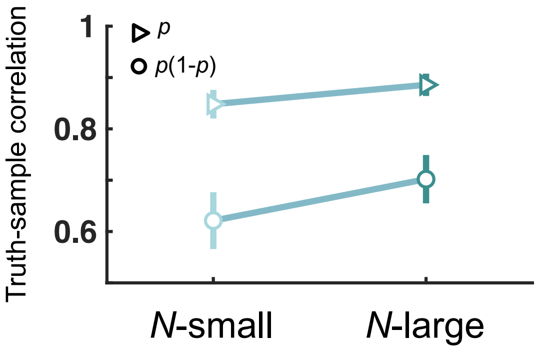


**S10 Fig. Consequences of random sampling errors.**

The *p* or *p*(1–*p*) estimated from a sample of dots from a display might deviate from the true value of *p* or *p*(1–*p*) in the display. As an evaluation of the fidelity of the sampled value, we used numerical simulations to compute the correlation (Pearson's *r*) between the true value and the sampled value, based on the BLO parameters estimated from individual subjects. Triangles and circles respectively denote *p* and *p*(1–*p*). The truth-sample correlation was higher in the *N*-large condition than in the *N*-small condition.
